# LRG1 is a novel HER3 ligand to promote growth in colorectal cancer

**DOI:** 10.1101/2023.02.18.529070

**Authors:** Moeez Rathore, Michel’le Wright, Daniel Martin, Wei Huang, Derek Taylor, Masaru Miyagi, Hao Feng, Yamu Li, Zhenghe Wang, Lee M Elis, Jordan Winter, Stephen Moss, John Greenwood, Rui Wang

## Abstract

Therapy failure for patients with metastatic colorectal cancer (mCRC) remains an overarching challenge in the clinic. We find that liver endothelial cells secrete soluble factor(s) to promote mCRC growth in vitro and in vivo. We identify LRG1 in ECs secretome, which promotes growth in tumor cells through binding and activation of HER3. Pharmacological blocking of the LRG1/HER3 axis using LRG1 antibody 15C4 completely attenuated LRG1-induced HER3 activation and in vitro and in vivo growth of the tumor. Moreover, LRG1-/- mice with CRC allografts in the liver had 2 times longer overall survival than tumor-bearing LRG1+/+ mice. Lastly, unbiased -omics analysis and target-specific inhibitors identified eIF4-protein synthesis is significantly activated by the LRG1/HER3/RSK1/2 axis. This work reveals a paracrine mechanism of mCRC growth in liver microenvironment and highlighted the potential of blocking LRG1-HER3 and involved downstream pathways for treating patients with mCRC.

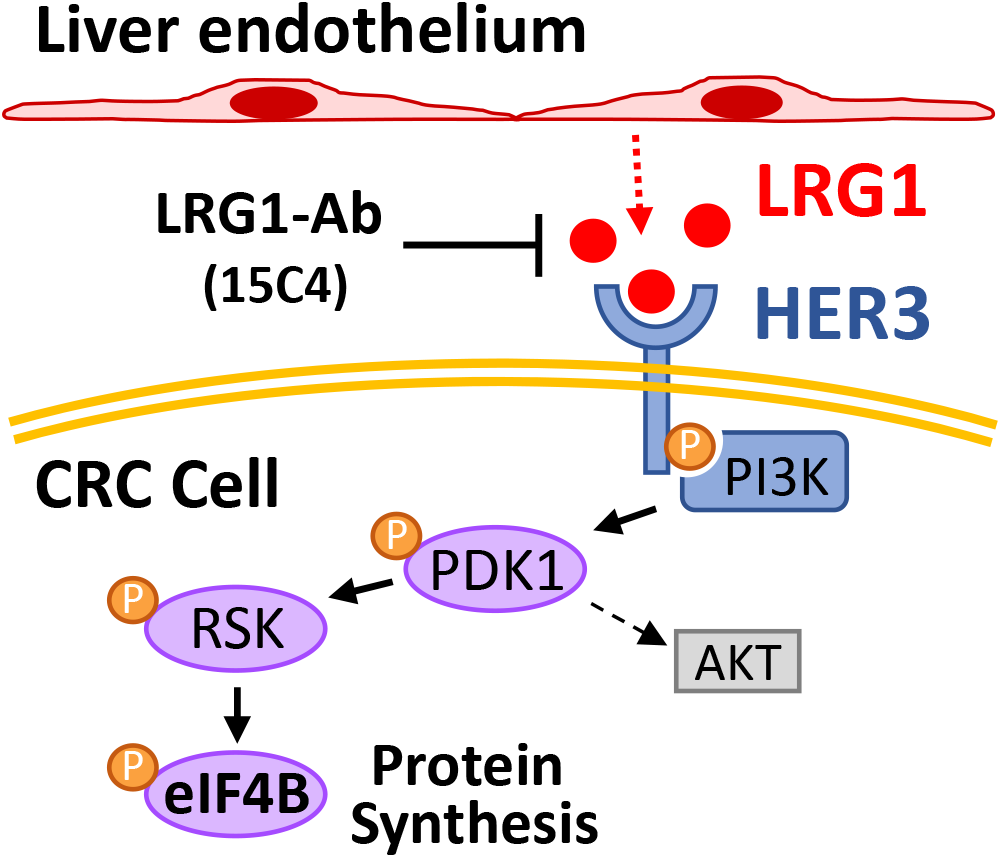

## INTRODUCTION

Colorectal cancer (CRC) remains the second leading cause of cancer-related death in the US ^1^. Patients with primary and localized CRC have 5-year survival rates between 53%-92% and are potentially curable by surgical resection and adjuvant therapy when appropriate ^2^. However, therapy failure for patients with metastatic CRC (mCRC) remains an overarching challenge as these patients only have a 5-year survival rate at 14%^1^, with the exception in patients with microsatellite instability-high tumors that benefit from the immune checkpoint blockade therapies ^3, 4^ (count for ∼5% of all mCRC ^5, 6^). Elucidating the regulation of survival pathways in mCRC is urgently needed for developing therapeutic interventions and improving outcomes for patients with mCRC.

Significant preclinical studies suggest human epidermal growth factor receptor 3 (HER3, also known as ErbB3) as a key factor for promoting growth and survival in CRC and other types of cancer ^7^. Upon binding to neuregulins, the only known ligand to date, HER3 activates AKT and other downstream pathways to promote cancer cell growth and survival ^8^. Most preclinical studies used subcutaneous (subQ) xenograft models and showed that blocking the HER3 signaling pathway attenuates tumor growth ^8, 9^. However, HER3-targeted therapies, all of which were developed based on the canonical neuregulin-HER3 interactions, led to limited impact in the clinic ^2, 7,10^. A better understanding of the regulatory mechanism of the HER3 pathway in clinically relevant animal models will help us to develop new strategies for blocking this pro-survival pathway in mCRC. In the context of CRC liver metastases that occur in ∼80% of all mCRC cases ^11^, we previously demonstrated that endothelial cells (ECs), a key component of the liver microenvironment, secret soluble factors to activate CRC-associated HER3 in a paracrine fashion that, in turn, promote CRC proliferation and survival ^12, 13^. We also showed that blocking HER3 significantly blocked mCRC tumor growth in an orthotopic model, suggesting HER3 activation by the liver microenvironment is a key in promoting mCRC development. Surprisingly, we found that the canonical HER3 ligand neuregulins are not involved in EC-induced HER3 activation, but the soluble factor(s) that activate HER3 have not been identified yet. Elucidating the potentially novel HER3 activation mechanism may lead to new strategies of blocking HER3-mediated mCRC survival.

In this study, we combined unbiased proteomics analyses with in-depth mechanistic studies and discovered that liver EC-secreted leucine-rich alpha-2-glycoprotein 1 (LRG1) is a novel HER3 ligand that promotes CRC cell growth. We determined that LRG1 inhibitions blocked mCRC tumor growth and prolonged mouse overall survival in syngeneic, orthotopic and proof-of-principle subQ models. We also identified the PI3K-PDK1-RSK-eIF4B signaling axis as a novel downstream target of LRG1-HER3 interactions, distinct from the canonical neuregulin-HER3 signaling. Overall, we determined that LRG1-HER3 signaling axis mediates the liver EC-cancer crosstalk for promoting mCRC development.

## RESULTS

### LRG1 is a new ligand that binds to and activates HER3

To further support the notion that HER3 activation is a key pro-survival pathway in mCRC, we report for the first time that HER3 activation, determined by phosphorylation, is higher in CRC liver metastases than in primary tumors or lymph node metastases (**Figure S1**), suggesting HER3 activation associates with mCRC development. We also used five distinct primary EC lines derived from human non-neoplastic liver tissues and validated that conditioned medium (CM) containing EC-secreted factors increased cell growth in CRC cells, including a PDX-derived cell line HCP-1 developed in prior studies ^14^ (**Figure S2A**), and that depleting the canonical HER3 ligand neuregulins from EC CM did no block EC-induced HER3 activation (**Figure S2B**).

To identify the liver EC-secreted soluble factor(s) that activate CRC-associated HER3, we first fractionated EC CM by fast protein liquid chromatography (FPLC) and found that soluble factors in fractions 7 and 8 activated HER3 CRC cells (**Figure 1A, B**). We then sent the remaining fractions 7 and 8 for protein identification by mass spectrometry (MS #1, **Table S1**). CRC CM were also processed and analyzed in parallel as controls. Separately, we used purified recombinant human HER3 extracellular domain (HER3 ECD) peptides to pull down the HER3-activating factor(s) from EC CM. As a result, EC CM after HER3 ECD pull-down failed to activate HER3, whereas the soluble factor(s) pulled down and eluted from HER3 ECD activated HER3 in CRC cells (**Figure 1C, D**). In parallel, Ni-NTA beads alone and HER2 ECD were used as negative controls, both failed to deplete HER3-activating factor(s) from EC CM. Moreover, EC CM depleted by HER3 ECD failed to promote cell growth, compared to complete EC CM and EC CM depleted by Ni-NTA beads or HER2 ECD controls (**Figure. 1E**). PDX-derived HCP-1 CRC cells were used for both MS studies. Similar effects of HER3 ECD depletion on EC CM were also observed when using CM from another primary EC line and with multiple CRC cell lines (**Figure S3A**). These data together suggest HER3 ECD sufficiently pulled down soluble factor(s) that bind and activate HER3. Subsequently, soluble factors eluted from HER3 ECD or controls (Ni-NTA beads and HER2 ECD) were subjected to MS analysis (MS #2, **Table S2**). CM of HCP-1 CRC cells were also processed for pull-down and MS assays in parallel as negative controls. Comparing the data from both MS analyses, we identified secreted soluble proteins that were exclusively found in EC CM fractions that activated HER3 (MS #1) and that were specifically bound to HER3 ECD (MS #2) (**Figure S3B**). Subsequent validations by treating CRC cells with identified candidate proteins revealed that leucine-rich alpha-2-glycoprotein 1 (LRG1) activated HER3 and promoted CRC cell growth (**Figure 1F**), whereas other assessed candidates did not activate HER3 (**Figure S3C**). We also determined that knocking down HER3 by siRNAs in CRC cells blocked LRG1-induced cell growth (**Figure S3D**), confirming LRG1-induced cell growth is mediated by HER3, which is supported our prior findings that HER3 inhibitions blocked EC CM-induced CRC cell and tumor growth ^12, 13^.

**Figure 1.**
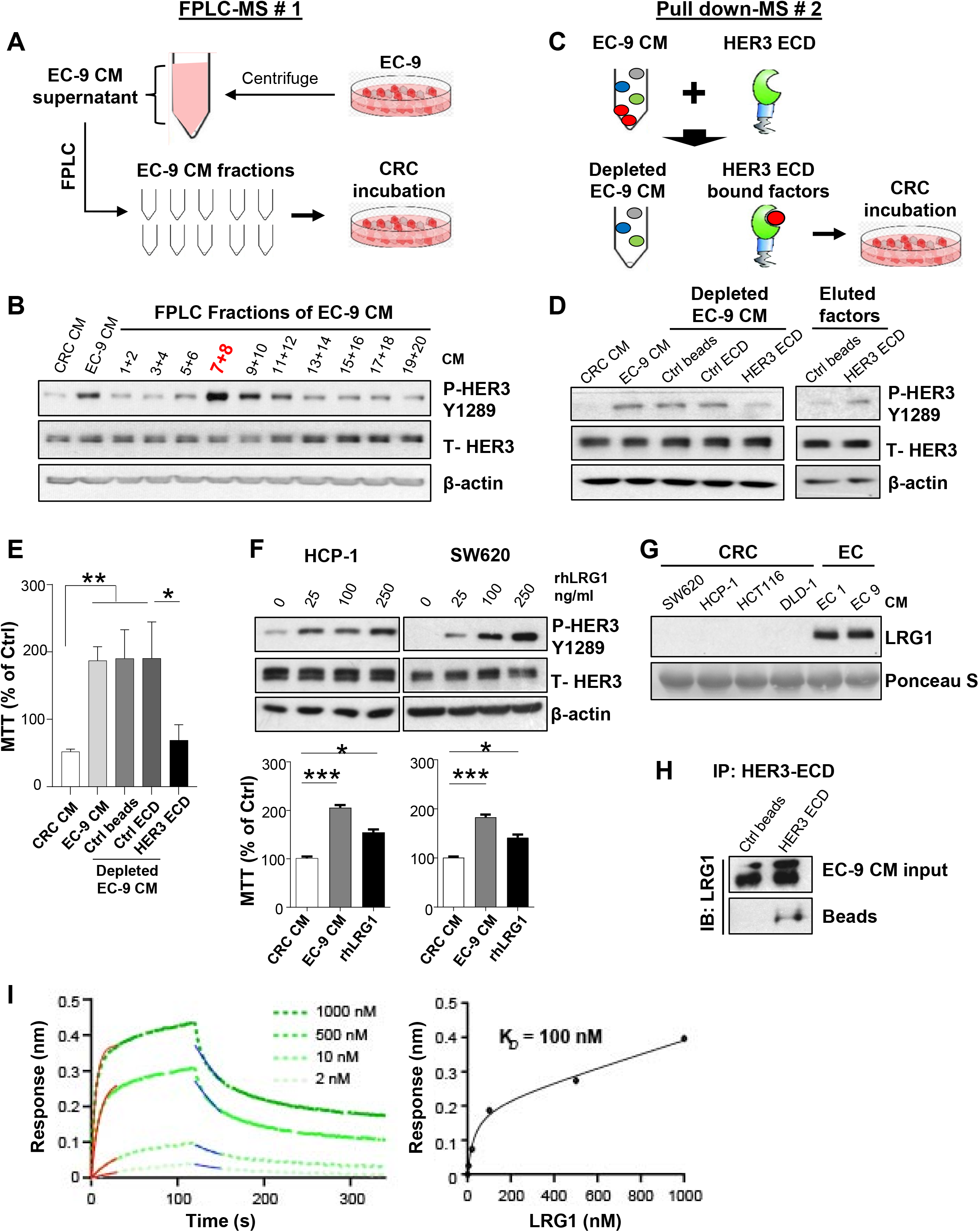
LRG1 binds to and activates HER3 as a novel ligand. **(A)** Schematics of CM fractionation by FPLC and subsequent treatment. **(B)** Western blotting showing phospho-HER3 (P-HER3) levels in HCP-1 CRC cells treated by CRC CM, complete EC-9 CM, or FPLC fractions of EC-9 CM. **(C)** Schematics of HER3 ECD pulling down soluble factors from EC-9 CM and subsequent treatment. Red dots indicate EC-secreted factor(s) that bind to HER3 ECD. **(D)** Western blotting showing P-HER3 levels in HCP-1 CRC cells treated by CRC CM, complete EC-9 CM, EC-9 CM after indicated depletions, or factors eluted from indicated beads or ECD. Ctrl beads, Ni-NAT beads. Ctrl ECD, HER2 ECD. **(E)** The MTT assay showing relative HCP-1 cell viability after treated by indicated CM. **(F)** Upper, Western blotting showing P-HER3 levels in CRC cells treated with recombinant human LRG1 (rhLRG1) at indicated levels. Lower, the MTT assay showing relative viability of CRC cells treated by indicated CM LRG1 (250 ng/ml). **(G)**. Western blotting showing soluble LRG1 in CM from CRC cells and ECs. **(H)** Western blotting showing LRG1 pulled down by control beads or HER3-ECD. **(I)** The binding affinity assay showing LRG1 and HER3-ECD binding affinity using LRG1 at indicated levels. K_D_=K_on_/K_off_ with 1:1 fit model. Total HER3 (T-HER3), β-actin, and Ponceau S are shown as loading controls. Data in (E) and (F) are shown as Mean +/-SEM, *p<0.05, **p<0.005, ***p<0.0005 by unpaired two-tailed *t*-test.

LRG1 is known to be secreted by ECs and activate the TGFβ receptor II -SMAD axis in ECs in autocrine fashion ^15^. We first confirmed that LRG1 was secreted by ECs but not CRC cells, and that LRG1 directly binds to HER3 as it was pulled down by HER3 ECD (**Figure 1G, H.** The canonical HER3 ligand neuregulins were not detected by HER3 ECD pull down). Meanwhile, HCP-1 and many established CRC cell lines harbor loss-of-function mutations in TGFβ receptors (CCLE database and published data ^16, 17^), thus, could not be activated by LRG1. We also showed that even in CRC cells with functional TGFβ receptors (SW620 and HT29), LRG1 activated HER3 without affecting SMAD3, a key readout of TGFβ receptor activation. Compared to that, TGFβ as a positive control activated SMAD3, as expected, but had no effect on HER3 (**Figure S3E**). Lastly, we performed a biolayer Interferometry (BLI) assay and determined the LRG1-HER3 binding affinity is at K_D_=100 nM (**Figure 1I**). In contrast, K_D_ of LRG1 for its known TGFβ receptor binding partners are over 200 nM ^15, 18^, suggesting lower binding affinity than LRG1-HER3. We also determined that LRG1-HER3 binding affinity is over three times higher than the canonical neuregulin-HER3 interaction (**Figure S3F**). Together, our convincing data determined that LRG1 directly binds to and activates HER3 as a novel ligand for promoting CRC cell growth, and that LRG1-HER3 binding is independent of the canonical LRG1-TGFβ receptor or neuregulin-HER3 interactions.

### LRG1 mediates EC-induced HER3 activation and growth in CRC cells

To further determine roles of EC-secreted LRG1 in HER3 activation and CRC cell functions, we first depleted LRG1 from EC CM by knocking down LRG1 in ECs with LRG1-specific siRNAs, with scramble siRNAs as a control (**Figure 2A**). We then treated CRC cells with complete or LRG1-depleted EC CM and determined that CM with LRG1 activated HER3 and increased CRC cell growth, as expected, whereas LRG1-depleted EC CM had no effect on HER3 activation or cell growth (**Figure 2B, C**). For additional validation, we depleted LRG1 from EC CM using a commercially available LRG1 antibody (**Figure 2D**), and showed that immuno-depletion of LRG1 also blocked HER3 activation and cell growth induced by EC CM (**Figure 2E, F**). To determine the effects of EC-secreted LRG1 on CRC tumor growth *in vivo*, we used a proof-of-principle subQ xenograft model developed in prior studies^12, 13, 19^. First, we subQ implanted CRC cell lines into athymic nude mice. Once xenografts were established, mice were randomized and then treated with CRC or EC CM by subQ injecting CM into the spaces between xenografts and skin tissues once a week. Compared to CRC CM controls, EC-9 CM led to significant 3-5-fold increases in tumor sizes. More importantly, immuno-depleting LRG1 from EC CM, as done in Figure 2D, completely abolished EC-induced xenograft growth (**Figure 2G**). Paired with that, IHC analysis of the xenografts showed that EC CM increased the levels of HER3 phosphorylation, indication of activation, which were decreased by LRG1 depletion (**Figure 2H**). Together, these findings from *in vitro* and *in vivo* studies demonstrated that LRG1 is the key mediator of EC-induced HER3 activation and CRC cell and tumor growth.

**Figure 2.**
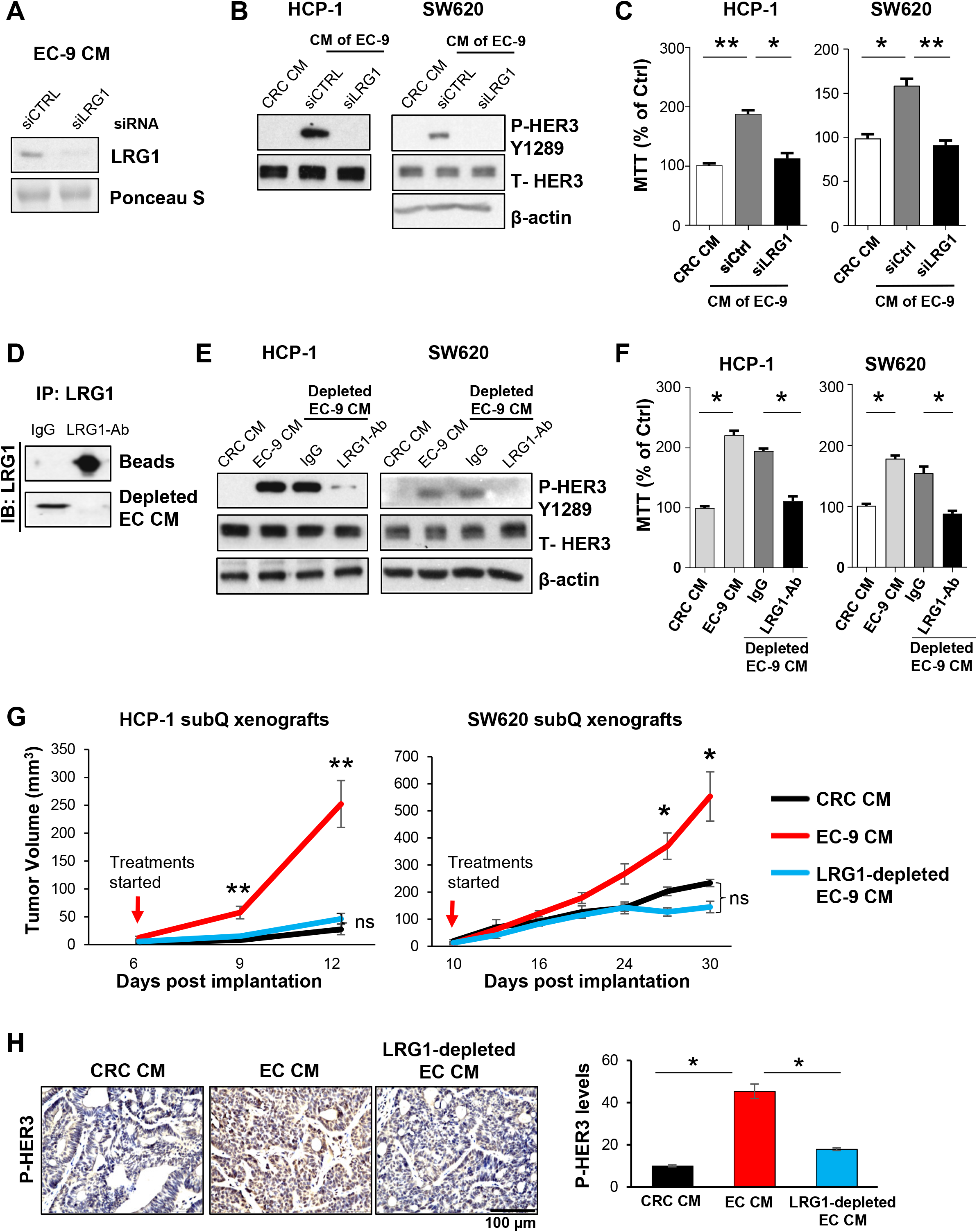
LRG1 blockades inhibit EC-induced HER3 activation and growth in CRC cells and xenografts. **(A)** Western blotting showing LRG1 levels in CM from ECs with control (siCTRL) or LRG1-specific siRNAs (siLRG1). **(B)** Western blotting showing P-HER3 levels in CRC cells treated by indicated CM. **(C)** The MTT assay showing relative viability of CRC cells treated by indicated CM. **(D)** Western blotting showing LRG1 pulled down by either control (IgG) or LRG1-specific (LRG1-Ab) antibodies and EC-9 CM after immuno-depletion. **(E)** Western blotting showing P-HER3 levels in CRC cells treated by CRC CM, complete EC-9 CM, or LRG1-depleted EC-9 CM. **(F)** The MTT assay showing relative viability in CRC cells treated by indicated CM. **(G)** Tumor volume changes of subQ xenografts with indicated treatments (n=10 mice/group). **(H)** Representative images of IHC staining of P-HER3 (Y1289) in HCP-1 xenografts and quantifications. Scale bar represents 100 µm. Ponceau S, total HER3 (T-HER3), and β-actin are shown as loading controls. Data in (C) and (F) are shown as Mean +/-SEM, *p<0.05, **p<0.005 by unpaired two-tailed *t*-test. Data in (G) and (H) are shown as mean -/+ SD, *p<0.05, **p<0.005 by one-way ANOVA *t*-test.

### LRG1 blockade increases mouse survival and inhibits mCRC tumor growth

To determine the effects of LRG1 on CRC liver metastases *in vivo*, we used mice with systemic LRG1 knockout (KO, *Lrg1*^-/-^) developed and described before^15^, and wild-type siblings (WT, *Lrg1*^+/+^), and used MC-38 murine CRC cells to develop CRC liver metastases in a syngeneic, orthotopic mCRC model. We found that tumor-bearing LRG1 KO mice had significantly longer medium overall survival than WT mice (72 days and 52 days, respectively. **Figure 3A**). Significant tumor burden in the liver was confirmed when mice became moribund and sacrificed (**Figure S4A**). We further determined the effects of blocking LRG1 on mCRC tumor growth using a LRG1 neutralizing antibody, 15C4, that was developed to neutralize human and murine LRG1 and demonstrated significant on-target effects in other disease models ^20–22^. After confirming 15C4 blocked EC-induced HER3 activation in a dose-dependent manner (**Figure S4B**), we first determined that 15C4 blocked EC-induced HER3 activation and cell growth in human and murine CRC cell lines *in vitro* (**Figure 3B, S4C**), and that 15C4 blocked the growth of subQ xenografts induced by LRG1-containing EC CM (**Figure 3C, S4D**). Interestingly, 15C4 had little effect on CRC cells or subQ xenografts without EC CM treatment, suggesting LRG1 blockade has limited effect on CRC primary tumors without the LRG1 stimulation from EC-rich microenvironment in the liver. Then, we assessed the effects of 15C4 in a syngeneic, orthotopic mCRC model in WT mice with LRG1 expression. We found that 15C4 phenocopied LRG1 knockout and prolonged mouse survival (70-day medium survival in 15C4-treated mice, 49-day in control mice, **Figure 3D**), and significantly decreased tumor growth (**Figure 3E**). Together, our findings determined that LRG1 plays a critical role in promoting mCRC growth, and that blocking LRG1 significantly blocks mCRC growth in the liver.

**Figure 3.**
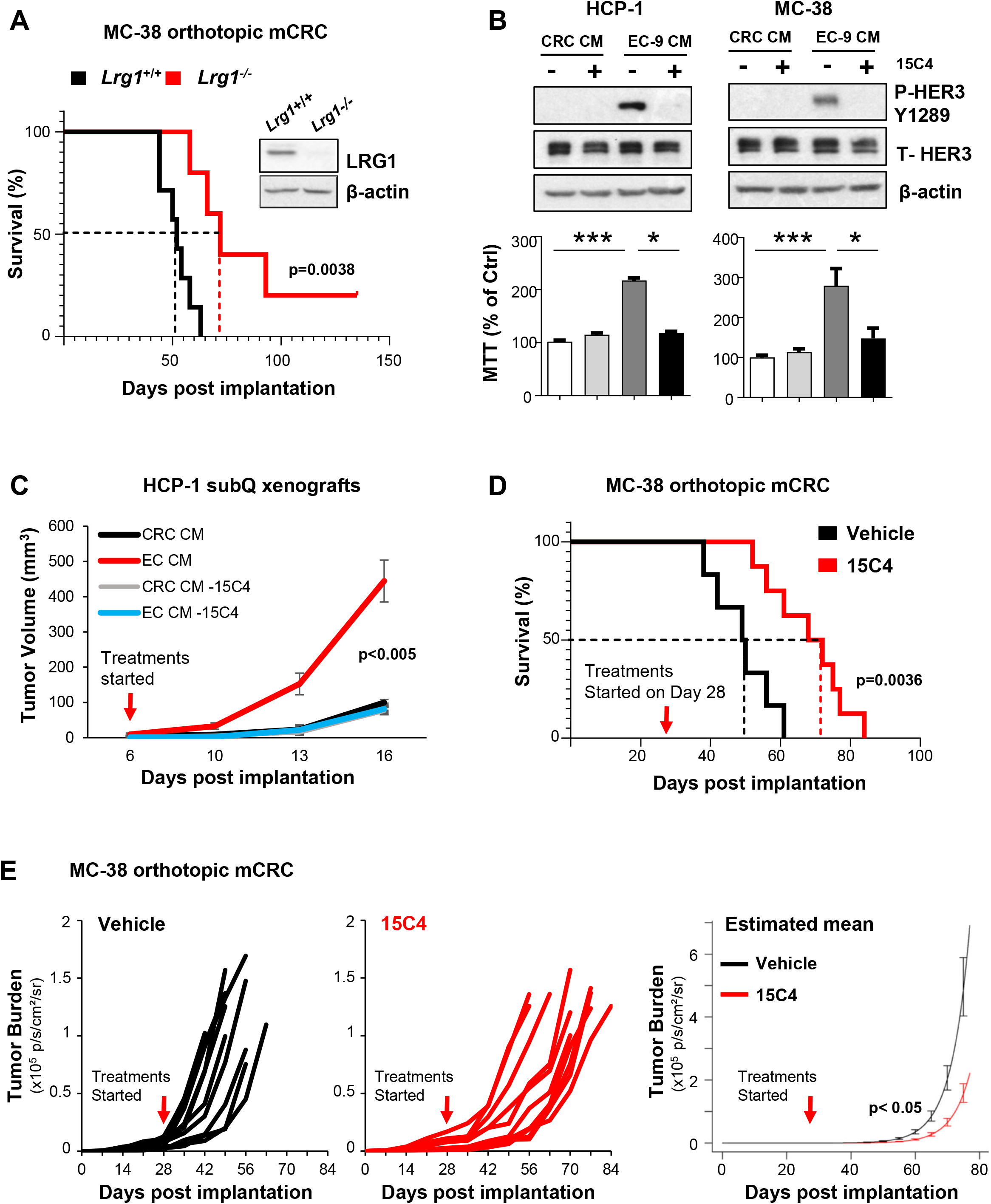
LRG1 blockades inhibit CRC tumor growth *in vivo*. **(A)** Kaplan-Meier curve showing overall survival of WT (*Lrg1*^+/+^) and LRG1 KO (*Lrg1^-/-^*) mice with CRC liver metastases (6 WT mice, 5 LRG1 KO mice), with Western blotting showing LRG1 levels in the liver. **(B)** Human (HCP-1) or murine (MC-38) CRC cells with indicated treatments of CM and 15C4 (500 µg/ml). Upper, Western blotting showing P-HER3 levels. Lower, the MTT assay showing relative cell viability. **(C)** Tumor volume of HCP-1 subQ xenografts with indicated treatments (5 mice/group). **(D)** Kaplan-Meier curve showing overall survival of mice with syngeneic, orthotopic mCRC treated with control vehicle or 15C4 (20 mg/kg, 10 mice/group). **(E)** Left and middle panels showing tumor burden of individual MC-38 orthotopic tumors with indicated treatments, right panel showing estimated mean tumor burden of each group. Total HER3 (T-HER3) and β-actin are shown as loading controls. P values in (A) and (D) are determined by Wilcoxon rank-sum test. Data in (B) are shown as Mean +/-SEM, *p<0.05, ***p<0.0005 by unpaired two-tailed *t*-test. Data in (C) and (E) are shown as Mean -/+ SD with p values determined by one-way ANOVA *t*-test.

### LRG1-HER3 axis activate eIF4B-mediated protein synthesis in CRC cells

To determine the downstream pathway(s) that are activated by the novel LRG1-HER3 interactions, we first compared the protein phosphorylation landscapes between CRC CM- and EC CM-treated CRC cells by global phospho-serine/threonine enrichment followed by LC-MS, and categorized the identified changes in protein phosphorylation into three groups: unique in CRC CM-treated cells, unique in EC CM-treated cells, and detected in both and significantly different between groups (**Figure 4A, Table S3**). We confirmed that CRC CM- and EC CM-treated groups are significantly different (**Figure S5A, B**), with the exception that one EC CM-treated sample had low correlation coefficient with other two EC CM-treated samples, thus, removed from further analysis. Subsequent QIAGEN Ingenuity Pathway Analysis (IPA) of the identified unique and significant changes revealed that protein synthesis is one of the top cellular functions and that eukaryotic initiation factor (EIF) signaling is the top pathway activated by EC CM (**Figure 4B—D, Table S3**). As EIF is a major mediator of eucaryotic cell protein synthesis^23^, our data strongly suggest that EC CM activate the EIF-protein synthesis signaling axis in CRC cells, which has not been reported in prior LRG1 or HER3 signaling studies. We validated our findings by assessing the effects of LRG1 on eIF4B, the top two identified EC CM-activated EIF components (**Figure 4E**), and a direct downstream target of receptor tyrosine kinases ^24^. Indeed, LRG1 increased eIF4B phosphorylation, indication of activation, in CRC cells (**Figure 4F**).

**Figure 4.**
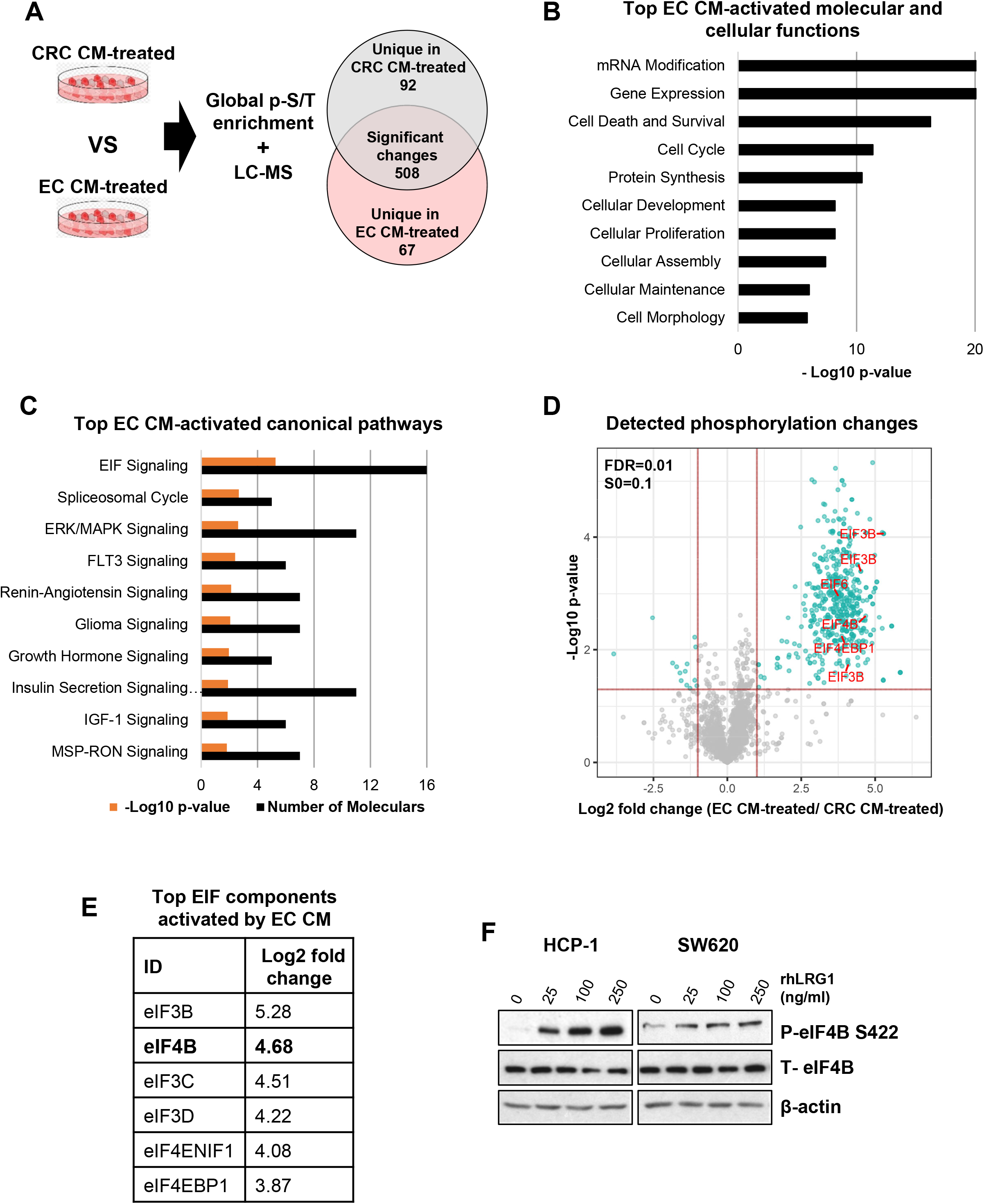
eIF4B is a novel downstream target of LRG1. **(A)** Schematics of global phospho-protein enrichment followed by LC-MS, with a pie chart showing distinct categories of identified changes. **(B)** Top 10 EC CM-activated molecular and cellular functions identified by IPA, ranked by -Log10 p values. **(C)** Top 10 EC CM-activated canonical pathways identified by IPA, ranked by -Log10 p values. **(D)** Volcano plot showing phospho-protein changes that are detected in both CRC CM and EC CM-treated groups. Blue dots highlight the 508 significant changes with > 2-fold change and p-value <0.05 by Student’s two-sample t-test. **(E)** Summary of IPA-identified top activated EIF components ranked by Log2 fold changes, also highlighted in red in panel (D). **(F)** Western blotting showing P-eIF4B levels in CRC cells treated by different levels of LRG1. Total eIF4B (T-eIF4B) and β-actin are shown as loading controls.

To further demonstrate that eIF4B-mediated protein synthesis is activated by the LRG1-HER3 interactions, we first used HER3-specific siRNAs and determined that HER3 knockdown blocked LRG1-induced eIF4B activation (**Figure 5A**). Coupled with that, the LRG1 neutralizing antibody 15C4 blocked LRG1-induced eIF4B activation and *de novo* protein synthesis in CRC cells (**Figure 5B, C**. Similar effects were observed when treating cells with EC CM). Moreover, we used eIF4 inhibitors (4EGI-1) and eIF4B-specific siRNAs, separately, and determined that eIF4B inhibitions completely blocked LRG1-induced protein synthesis and growth in CRC cells (**Figure 5D, S6A—C**). Lastly, we assessed subQ xenografts and orthotopic CRC liver metastases from studies showed in Figures 2G and 3D, and found that LRG1 inhibitions blocked EC-induced eIF4B activation in CRC tumors *in vivo* (**Figure 5E, F**). Of note, we found that LRG1 inhibition had no effect on other eIF4 subunits (**Figure S6D** show data for eIF4G as an example). Together, these findings identify eIF4B-mediated protein synthesis as a novel downstream pathway of the LRG1-HER3 axis, and suggest that eIF4B mediates LRG1-HER3 induced CRC growth.

**Figure 5.**
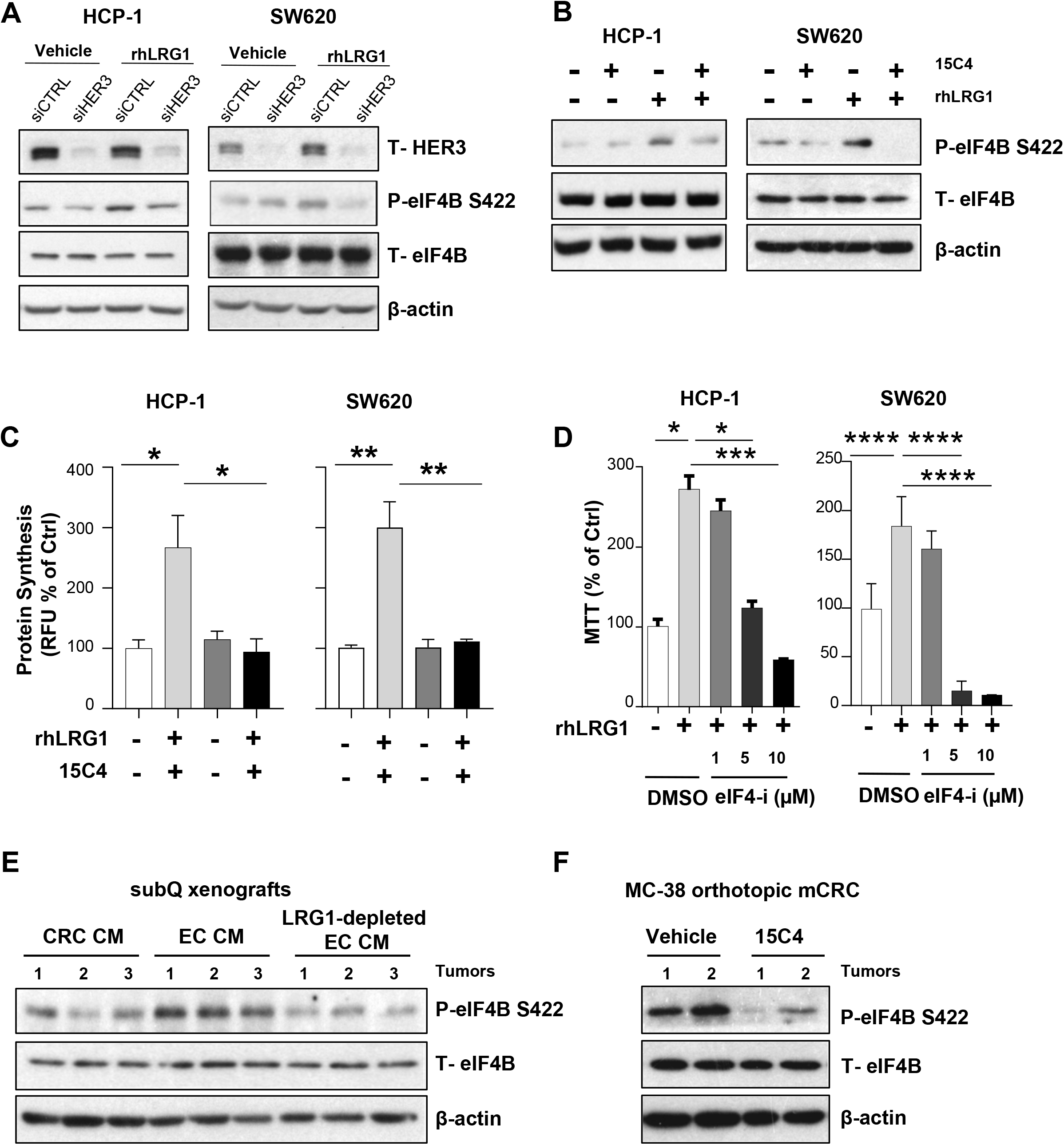
The LRG1-HER3 axis activates eIF4B to promote protein synthesis and growth in CRC cells. **(A)** Western blotting showing P-eIF4B levels in CRC with control (siCTRL) or HER3-specific siRNAs (siHER3) and treated with LRG1. **(B)** Western blotting showing P-eIF4B levels in CRC cells treated by LRG1 and 15C4. **(C)** The protein synthesis assay showing relative levels of synthesized *de novo* proteins in CRC cells treated with/without LRG1 and 15C4 as indicated. **(D)** The MTT assay showing relative viability in CRC cells treated with/without LRG1 and eIF4 inhibitors 4EGI-1 as indicted. (**E, F**). Western blotting showing P-eIF4B levels in individual CRC subQ xenografts and orthotopic tumors after indicated treatments. 250 ng/ml LRG1 and 500 µg/ml 15C4 were used for all *in vitro* assays. Total HER3 (T-HER3), total eIF4B (T-eIF4B), and β-actin are shown as loading controls. Data in (C) and (D) are shown as Mean +/-SEM, *p<0.05, **p<0.005, ***p<0.0005, ****p<0.00005 by unpaired two-tailed *t*-test.

### The PI3K-PDK1-RSK signaling axis mediates LRG1-induced eIF4B activation

To elucidate the specific mechanism(s) by which LRG1-HER3 interactions activate eIF4B, we first compared the effects of LRG1 and the canonical HER3 ligand neuregulins on CRC cells and found that although HER3 was activated by both proteins, eIF4B was only activated by LRG1 but not neuregulins (**Figure. 6A.** Additional validations are shown in **Figure S7A**). We also assessed the effects of LRG1 and neuregulins on AKT as it is an established upstream effector for eIF4B activation^25^, and is a downstream target of HER3 by canonical neuregulin binding^7^. Neuregulins activated AKT, as expected, but LRG1 had no effect on AKT (**Figure 6A**). To further determine that AKT is not involved in LRG1-induced CRC cell functions, we used an allosteric AKT inhibitor, MK-2206, and found that blocking AKT had no effect on LRG1-induced eIF4B activation (**Figure S7B**). Meanwhile, prior studies reported that 90 kDa ribosomal S6 kinase (RSK) family proteins can also activate eIF4B by direct phosphorylating the S422 residue we assessed ^24–26^, and that RSK can be activated independent of AKT by the IRS1-PI3K-PDK1 axis ^27–29^. Therefore, assessed the potential of LRG1-HER3 interactions recruiting the IRS1-PI3K-PDK1 axis to activate RSK, which then activating eIF4B in CRC cells. Indeed, we found that RSK was only activated by LRG1 but not neuregulins, determined by RSK1 phosphorylation at multiple sites (**Figure. 6B**). Then, we used specific inhibitors against IRS (by NT157), PI3K (by BYL719, a p110α inhibitor), PDK1 (by BX795), or RSK (by BI-D1870, a pan-RSK inhibitor) separately and determined that blocking the IRS-PI3K-PDK1 axis attenuated LRG1-induced eIF4B activation and protein synthesis (**Figure 6C, D**). In contrast, AKT inhibitors had no effect on LRG1-induced protein synthesis, further validated that AKT is not involved in LRG1-indcued CRC cell functions. We also used inhibitors against p110β as it is another PI3K isoform expressed in CRC cells, and validated that blocking p110β inhibitor blocked LRG1-induced eIF4B activation (**Figure S7C**). Moreover, we further validated our findings in HCT116 cells (**Figure S8**), another CRC cell line that does not have functional TGFβ receptor II for canonical LRG1 signaling (CCLE database). Together, our findings demonstrated that LRG1-HER3 interactions activate the IRS-PI3K-PDK1-RSK signaling cascade to induce eIF4B-mediated protein synthesis in CRC cells (model summarized in **Figure 6E**). Our data also determined that the LRG1-induced signaling cascade is distinct from the canonical neuregulin-HER3 interactions and independent of AKT.

**Figure 6.**
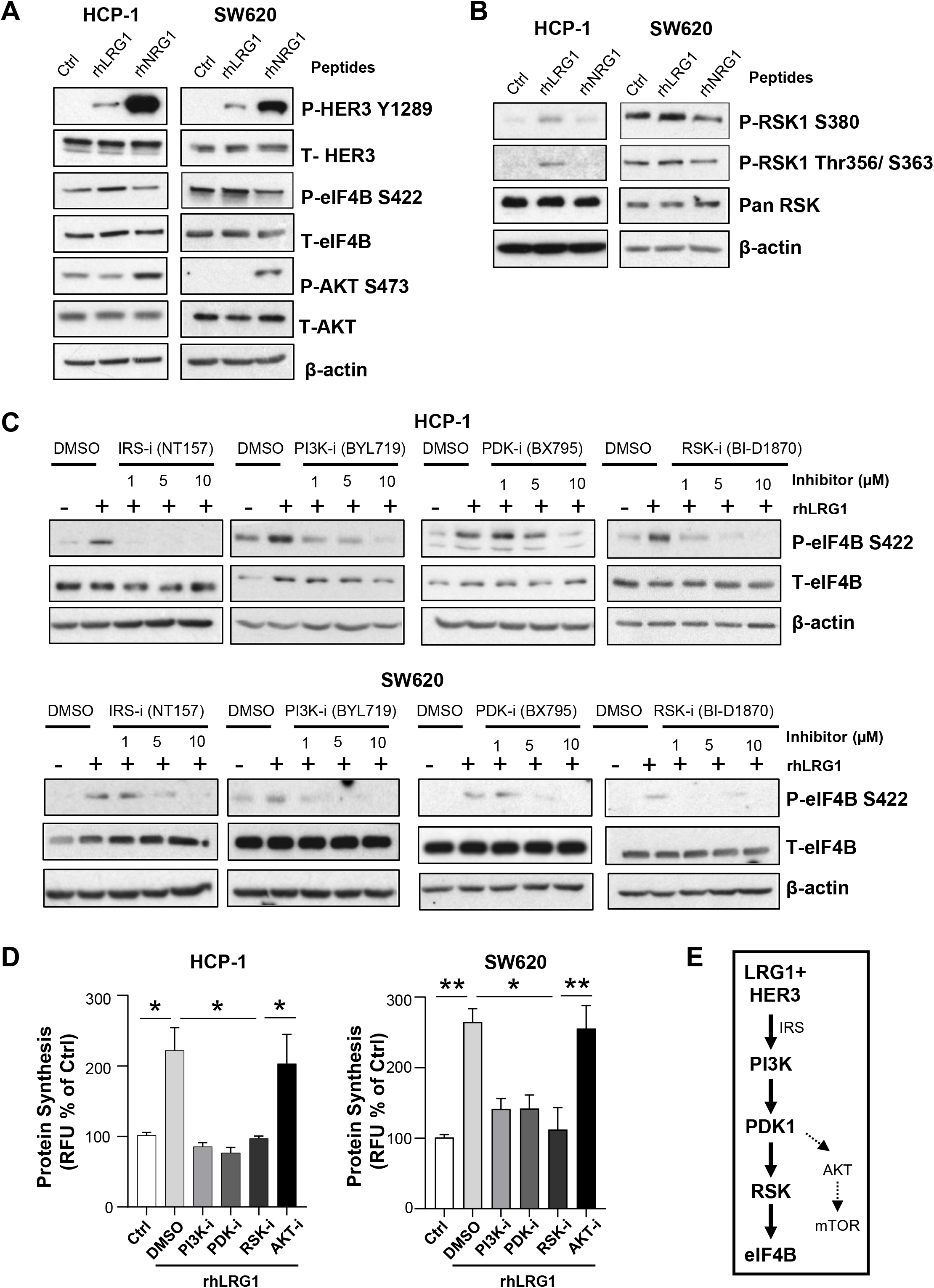
The PI3K-PDK1-RSK axis mediates LRG1-induced eIF4B activation and protein synthesis in CRC cells. **(A, B)** Western blotting showing P-HER3, P-eIF4B, P-AKT, and P-RSK1 levels in CRC cells treated with LRG1 or neuregulins (NRG1). **(C)** Western blotting showed P-eIF4B levels in CRC cells treated with LRG1 and indicated levels of inhibitors for IRS (NT157), p110α (BYL719), PDK1 (BX795), and RSK (BI-D1870), with DMSO as solvent controls. **(D)** The protein synthesis assay showing relative levels of synthesized *de novo* proteins in CRC cells treated with/without LRG1 and indicated specific inhibitors. **(E)** Schematics of the determined signaling pathway. Total HER3 (T-HER3), total eIF4B (T-eIF4B), total AKT (T-AKT), total RSK (pan RSK), and β-actin in (A-C) are shown as loading controls. Data in (D) are shown as Mean +/-SEM, *p<0.05, **p<0.005 by unpaired two-tailed *t*-test.

### RSK1 and RSK2 mediates LRG1-induced eIF4B activation

We first assessed RSK1 as it is a direct downstream target of the PI3K-PDK1 axis^30^. Since CRC cells express RSK isoforms 1, 2, and 3 but not RSK4 ^31^, we aimed to further determine the effects of LRG1-HER3 interactions on RSK isoforms 1-3 and identify the specific RSK isoform(s) that mediate LRG1-induced eIF4B activation. First, we confirmed that RSK1 is consistently activated by either EC CM or LRG1 peptides in multiple CRC cell lines, and determined that depleting LRG1 from EC CM or knocking down HER3 in CRC cells blocked LRG1-induced RSK1 activation (**Figure 7A, B**). In contrast, LRG1 had no effect on RSK2, and activated RSK3 in HCP-1 but not in other CRC cells (**Figure S9.** SW620 and HCT116 had no detectable RSK3 phosphorylation). Then, we used siRNAs to knock down RSK isoforms. Interesting, we found that knocking down either RSK1 or RSK2 alone led to elevated expression of the other RSK isoform in CRC cells (**Figure 7C**). Coupled with that, knockdown of RSK1 or -2 alone had no effect on eIF4B activation, whereas dual knockdown of RSK1 and -2 blocked LRG1-idncued eIF4B activation (**Figure 7D**). These findings suggest that RSK1 and -2 compensate each other when blocking one isoform and have redundant effects on eIF4B activation, which were observed in other types of cancer ^26, 30^. On the other hand, RSK3 siRNAs only modestly decreased eIF4B activation in one cell line and combinations with RSK1 or -2 siRNAs did not block eIF4B activation. Our data suggest that LRG1-HER3 interactions specifically activated RSK1. However, as CRC cells have significant RSK2 basal activity and that RSK1 and -2 have redundant effects on downstream targets, blocking both isoforms are necessary to sufficiently block LRG1-induced eIF4B activation and cell functions, as also demonstrated by findings with pan-RSK inhibitors in Figure 6.

**Figure 7.**
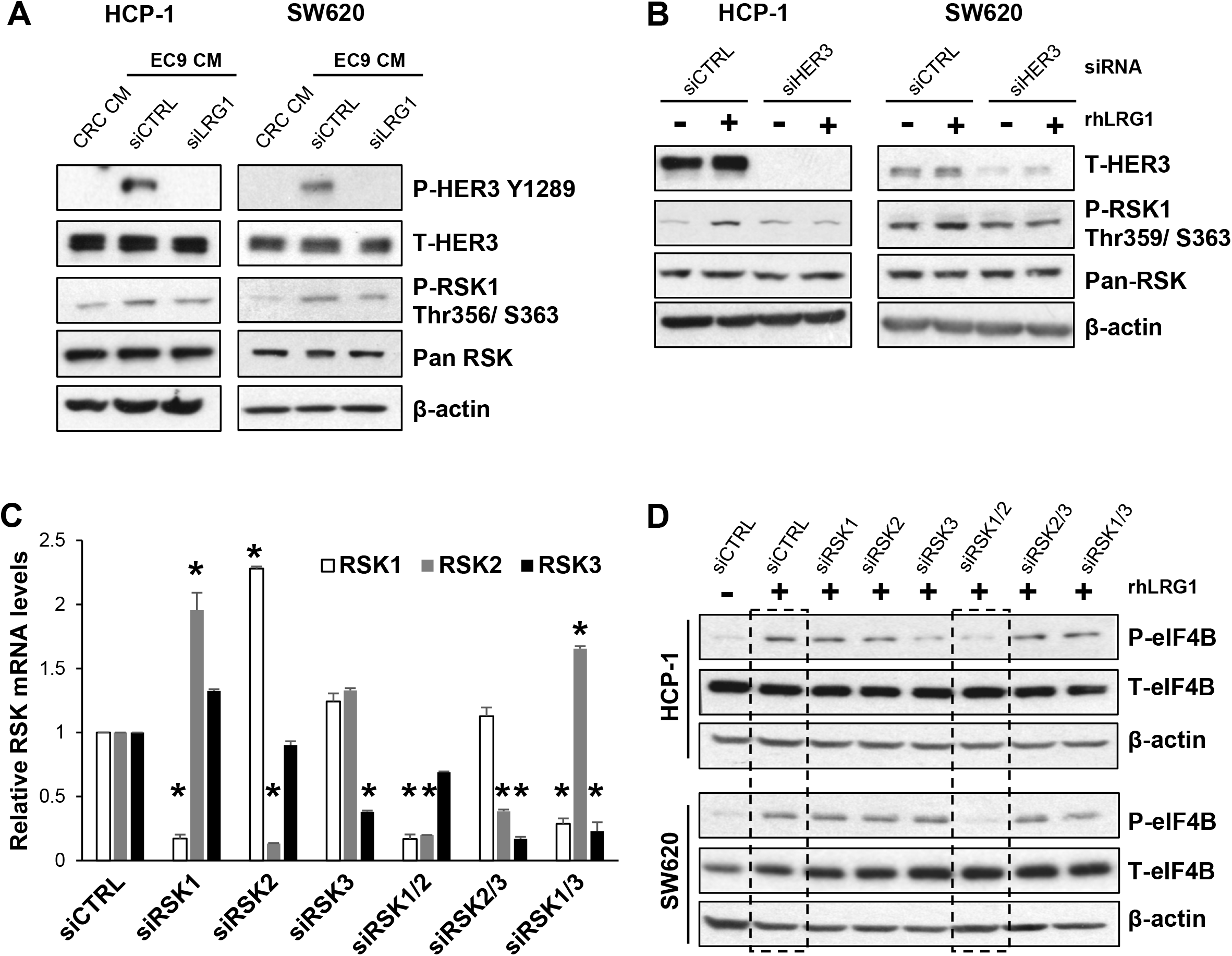
LRG1-HER3 activates RKS1 and RSK1/2 dual inhibition blocks LRG1-induced eIF4B activation. **(A)** Western botting showing P-HER3 and P-RSK1 levels in CRC cells treated by CRC CM and CM from EC-9 with control (siCTRL) or LRG1-specific siRNAs (siLRG1). **(B)** Western blotting showing total HER3 (T-HER3) and P-RSK1 levels in CRC cells with control (siCTRL) or HER3-specific siRNAs (siHER3) and treated with LRG1. **(C)** mRNA levels of RSK1, 2, and 3 in CRC cells with indicated siRNAs, determined by qPCR. **(D)** Western blotting showing P-eIF4B levels in CRC cells treated by LRG1 and indicated RSK siRNAs. Dash line boxes highlight eIF4B activation by LRG1 and inhibition by RSK-1 and -2 siRNAs, respectively, for direct comparison. Total HER3 (T-HER3), total RSK (pan RSK), total eIF4B (T-eIF4B) and β-actin are shown as loading controls. Data in (C) are also shown as Mean +/-SEM, *p<0.05 by unpaired two-tailed *t*-test compared to control siRNA (siCTRL) groups.

## DISCUSSION

CRC distant metastases are predominantly found in the liver, which has a unique EC-rich microenvironment with up to 20% of the liver parenchyma composed of liver ECs ^32–34^. Unlike precedent EC studies focused on angiogenesis and vascular formation, we determined that ECs secrete soluble factors to promote mCRC development in a paracrine fashion. Leveraging primary human liver ECs established in our laboratory, we discovered that LRG1 secreted from liver ECs activates CRC-associated HER3 as a novel ligand and promotes CRC cell growth, and the ligand-receptor interaction is independent of canonical LRG1 and HER3 pathways described previously. The significant pro-survival role of the LRG1-HER3 signaling axis was first demonstrated by *in vitro* studies showing LRG1-containg EC CM and purified LRG1 peptides activate HER3 and promote CRC cell growth, whereas depleting LRG1 from EC CM or blocking LRG1 with the neutralizing antibody 15C4 completely blocked EC-induced HER3 activation and cell growth. By performing four distinct animal studies either in a proof-of-principle subQ xenograft model or in a syngeneic, orthotopic model with CRC liver metastasis, we showed collectively that EC-secreted LRG1 promotes CRC tumor growth, and that blocking LRG1 either by immuno-depletion, LRG1 gene knockout, or the LRG1-specific antibody 15C4 blocked tumor growth and prolonged overall survival in mCRC-bearing mice. Moreover, we identified eIF4B-mediated protein synthesis as a novel downstream pathway of LRG1-HER3, distinct from the canonical neureguling-HER3 interaction, and determined that the PI3K-PDK1-RKS signaling cascade mediates LRG1-induced eIF4B activation independent of AKT (**Figure 6E**). Altogether, we identified LRG1 as a novel HER3 ligand and determined LRG1-HER3 as a key pro-survival signaling pathway that mediates mCRC growth triggered by the liver EC-CRC crosstalk.

LRG1 was first discovered to be expressed and secreted by retinal ECs as an autocrine factor that activates TGFβ receptor II and downstream SMAD pathway for promoting angiogenesis in eye diseases ^15^. Recent studies reported that LRG1 is also expressed in melanoma and pancreatic cancer and activates EGFR for promoting cancer cell growth ^35, 36^. They showed that knocking down LRG1 in cancer cells decreased cell and tumor growth, whereas knocking out LRG1 in host animals had no effect on B16F10 melanoma tumors, suggesting intracellular LRG1 is the key for regulating cancer cell functions. Distinct from those studies, we determined that LRG1 does not activate the TGFβ receptor-SMAD signaling axis in CRC cells (**Figure S3E**), and we previously reported that EGFR is not involved in EC-induced HER3 activation ^12, 13^. Instead, we showed that LRG1 directly binds to HER3 with high affinity and activates HER3 to promote CRC cell growth. B16F10 cells have low HER3 expression ^37^, which may explain why LRG1 knockout in host animals had little effect on B16F10 tumors. Also, in those studies, B16F10 tumors were implanted subQ, of which the surrounding microenvironment does not have the EC-rich niche in the liver for providing significant LRG1 stimulation. Indeed, we also found that LRG1 inhibitions had no effect on CRC subQ xenografts without stimulation, but completely blocked subQ tumor growth induced by LRG1-containing EC CM (**Figure 2G, 3C**). Our findings not only identified a novel mechanism of LRG1 activating HER3 for promoting cancer cell growth, but also suggest that the pro-cancer role of LRG1 is more significant in the presence of liver ECs with significant LRG1 stimulation. We focused on the liver EC microenvironment in the context of CRC liver metastases. Interestingly, LRG1 is also secreted by hepatocytes ^38, 39^, another key component of the liver microenvironment. Although hepatocyte is not the focus of this project and was not assess, it suggests that liver ECs and hepatocytes together provide significant levels of LRG1 in the liver that activates the pro-survival HER3 signaling in CRC liver metastases. Together, our findings described a paradigm-shifting discovery that extracellular soluble LRG1 in the liver activates CRC-associated HER3 as a new ligand and promotes CRC cell growth. The significant anti-cancer effects of LRG1 blockades demonstrated in orthotopic mCRC models highlight a potential strategy of blocking the liver-cancer crosstalk mediated by LRG1-HER3 for treating patients with CRC liver metastases.

LRG1 expression and the functions of intracellular LRG1 in CRC cells remain unclear. On one hand, our data (**Figure 1G**), and others^40^, show that CRC cells do not express LRG1. On the other hand, a separate group detected intracellular LRG1 in human CRC cells and tissues using a LRG1 antibody different from the one we used ^41–43^. However, they failed to detect the decreased LRG1 protein levels caused by siRNAs, and reported that LRG1 knockdown had no significant effect on CRC cell growth or other functions, suggesting LRG1 may not be expressed in CRC cells and intracellular LRG1 had little effect on cancer cell functions. Interesting, a recent study detected LRG1 in intestinal polyps and tumors development in GEMM models and showed that systemic knockout of LRG1 decreased intestinal polyp and tumor growth ^22^. However, they suggested that the effects on polyps and tumors were not caused by cancer-derived LRG1, instead, were driven by altered LRG1 signaling in the vasculature without changing vessel density. Coupled with the findings that LRG1 also promote Th17 inflammation and other pro-cancer signaling pathways ^39, 44^, roles of LRG1 in the tumor microenvironment and cancer intracellular signaling in CRC need to be further elucidated in future studies.

The canonical neuregulin-HER3 interactions require HER3 dimerization with other HER family receptors (EGFR, HER2 and HER4), which then leads to HER3 trans-phosphorylation and activation of downstream survival pathways ^7^. We previously showed that blocking other HER receptors had no effect on the neuregulin-independent HER3 activation induced by EC-secreted factors ^12, 13^. In the present study, we used purified LRG1 and HER3 peptides and further determined that LRG1 directly binds to HER3 without other HER receptors. Moreover, we directly compared and determined that the binding affinity of LRG1-HER3 is over 3 times higher than that of neuregulin-HER3 (**Figure S3F**). Although it is not clear whether HER3 dimerizes with another protein after LRG1 binding, our data strongly suggest that LRG1 and neuregulins have distinct binding mechanisms with HER3, which may explain why neuregulins were not detected in pull down assays with purified HER3 peptides. The differences between LRG1 and neuregulins effects on HER3 and CRC cell functions are further highlighted by the findings that the PI3K-PDK1-RSK-eIF4B signaling cascade is specifically activated by LRG1 but not neuregulins. Although the PI3K-PDK1-RSK signaling and RSK activating eIF4B have been reported separately in prior studies ^24–29^, we identified RSK-eIF4B as a novel downstream target of HER3 and determined for the first time that the PI3K-PDK1 signaling activates eIF4B independent of AKT. Moreover, our data suggest that although AKT is highly active in CRC cells, AKT inhibition have little effect on CRC cells in the context of LRG1-HER3-induced eIF4B-protein synthesis and survival, as shown by our data with AKT inhibitors.

HER3-targeted therapies developed to date are based on the canonical neuregulin-HER3 binding and have made little impact in the clinic ^7^, with the exception in patients with rare *NRG1* fusion mutations ^45–48^. As we determined that LRG1-HER3 interactions are distinct from the canonical neuregulin-HER3 binding, HER3-targeted therapies may not block the novel pro-survival LRG1-HER3 signaling cascade. Our findings suggest LRG1 inhibitions as a novel approach for blocking LRG1-HER3 signaling in mCRC, and potentially other types of cancer that express HER3. We also identify the downstream RSK-eIF4B signaling as an amenable target, of which specific inhibitors have been assessed in human trials ^49, 50^. Lastly, further characterizing the LRG1-HER3 binding mechanism will help to develop effective approaches to block the pro-cancer LRG1-HER3 interactions.

## ACKNOWLEDGMENTS

We thank Dr. Belinda Willard, Director of the CCCC Proteomics and Metabolomics Core for conducting global phospho-serine/threonine enrichment followed by LC-MS analysis. This project was supported by the NIH grant CA225756 (R.W.) and NIH institutional grants P30CA043703 (CWRU CCSG).

## AUTHOR CONTRIBUTIONS

Conceptualization, M.R. and R.W.; Methodology, M.R., D.T., J.W., Z.W., and R.W.; Validation, M.R., M.W., M.M., and R.W.; Formal Analysis, M.R., D.M., M.M., H.F., M.R., and R.W.; Investigation, M.R., M.W., W.H, M.M., Y.L.; Resources, Z.W., L.M.E., S.M., and J.G.; Data Curation, D.M., M.M., H.F., and R.W.; Writing – Original Draft, M.R. and R.W.; Visualization, M.R. D.M., W.H., M.M., H.F., and R.W.; Supervision, D.T., Z.W., J.W., S.M., J.G., and R.W.; Funding Acquisition, R.W.;

## DECLARATION OF INTERESTS

J.G. and S.E.M. are founders of a company spun out by UCL Business to commercialize a LRG1 function-blocking therapeutic antibody developed through the UK Medical Research Council DPFS funding scheme. This is currently directed toward treating ocular disease. J.G. and S.E.M. are members of the scientific advisory board and are shareholders of this company and named inventors on three patents related to LRG1 as a therapeutic target.

## SUPPLEMENTAL INFORMATION TITLES AND LEGENDS

**Figure S1. Human mCRC liver metastases have high levels of HER3 phosphorylation.** Representative images of IHC staining and quantifications showing phospho-HER3 (P-HER3) at Y1289 in colon mucosa and indicated CRC tissues from different stages. Sections were counter stained with hematoxylin. Scale bars 100 µm. Mean +/-SD, *p<0.05, **p<0.005 by one-way ANOVA test.

**Figure S2. CM form primary liver ECs increase CRC cell growth and HER3 activation independent of NRGs. (A)** The MTT assay showing relative cell viability in CRC cells treated by conditioned medium (CM) from each CRC cell line or from different EC lines. **(B)** Western blotting showing levels of neuregulins (NRG) in CM from EC-9 cells with control (siCTRL) or neuregulin-specific siRNAs (siNRG1-1 and -2. Other NRG isoforms not expressed in ECs), and P-HER3 levels in CRC cells treated by CRC CM, or CM from EC-9 with indicated siRNAs. Ponceau S, total HER3 (T-HER3), and β-actin are shown as loading controls. Data in (A) are shown as Mean +/-SEM, **p<0.005, ***p<0.0005 by unpaired two-tailed *t*-test.

**Figure S3. Analysis of candidate proteins identified by MS assays using EC CM fractions and HER3 ECD pull-down, related to** Figure 1**. (A)** Western blotting showing P-HER3 levels in CRC cells treated by CRC CM, complete EC-1 CM or EC-1 CM with indicated depletions. **(B)** Pie charts to demonstrate the numbers of proteins in each category identified by MS # 1 and # 2. **(C)** A list of top candidate proteins identified by MS analysis (ranked by intensity determined in MS # 2), and Western blotting showing P-HER3 levels in CRC cells treated by indicated proteins. Five top identified candidate proteins were validated (highlighted in black). recombinant human NPR1 and LTBP3 (in grey) are not available and not assessed. **(D)** The MTT assay showing relative cell viability in CRC cells with control (siCTRL) or HER3-specific (siHER3) siRNAs and treated with LRG1 (250 ng/ml). **(E)** Western blotting showing P-HER3 and P-SMAD3 levels in CRC cells treated by LRG1 or TGFβ peptides at 250 ng/ml. Both SW620 and HT29 cells have functional TGFβ receptors. **(F)** The binding affinity assay showing estimated EC50 of HER3 ECD binding with either LRG1 or neuregulins (NRG1). Total HER3 (T-HER3), total Smat3 (T-Smad3), and β-actin are shown as loading controls. Data in (D) are shown as mean +/-SEM, **p<0.005 by unpaired two-tailed *t*-test.

**Figure S4. LRG1 blockades delay CRC tumor growth and inhibit HER3 activation, related to** Figure 3**. (A)** Representative pictures of tumor-bearing livers collected from WT (*Lrg1*^+/+^) and LRG1 KO (*Lrg1^-/-^*) mice when mice became moribund and sacrificed. Pale light regions are CRC tumors and dark red regions are non-neoplastic liver parenchyma. Scale bars represent 1 cm. **(B)** Western blotting showing P-HER3 levels in CRC cells treated by indicated CM and the LRG1 neutralizing antibody 15C4 at different levels. **(C)** Human CRC cells treated with indicated CM and 15C4 (500 µg/ml). Left, Western blotting showing P-HER3 levels; right, the MTT assay showing relative cell viability. **(D)** Weights of HCP-1 subQ xenografts from mice treated with CM and 15C4 (n=5 mice/group). Total HER3 (T-HER3) and β-actin are shown as loading controls. Data in (C) are shown as Mean +/-SEM, ***p<0.0005 by unpaired two-tailed *t*-test. Data in (D) are shown as Mean -/+ SD, *p<0.05 by one-way ANOVA *t*-test.

**Figure S5. Distinct protein phosphorylation profiles in CRC CM- and EC CM-treated CRC cells, related to** Figure 4**. (A)** Scattered plot comparing phospho-protein changes detected in three replicates of CRC CM or EC CM-treated CRC cells, with Pearson correlation coefficients highlighted in blue. Coefficients >0.94 indicate high correlation. **(B)** Principal component analysis clustering CRC CM- and EC CM-treated samples with variances of 75.2% and 11.7% between clusters.

**Figure S6. eIF4B knockdown blocks LRG1-idnuced protein synthesis and growth in CRC cells, related to** Figure 5**. (A)** Western blotting showing eIF4B levels in CRC cells transfected with control (siCTRL) or eIF4B-specific (si-eIF4B) siRNAs. **(B)** The protein synthesis assay showing relative levels of synthesized *de novo* proteins in CRC cells with indicated treatment of eIF4B siRNAs and LRG1. **(C)** The MTT assay showing relative viability of CRC cells with indicated treatments. **(D)** Western blotting showing P-eIF4G in CRC cells treated by CRC CM, complete EC CM, or LRG-depleted EC CM. β-actin and total eIF4GB (T-eIF4G) are shown as loading controls. Data in (B) and (C) are shown as Mean +/-SEM, *p<0.05, **p<0.005 by unpaired two-tailed *t*-test.

**Figure S7. eIF4B is activated by LRG1 and blocked by p110**β **inhibitors, but not affected by AKT inhibitors, related to** Figure 6**. (A)** Western blotting from additional replications showing levels of P-HER3 and P-eIF4B in CRC cells treated by LRG1 and neuregulins (NRG1). **(B)** Western blotting showing P-AKT and P-eIFB4 levels in CRC cells treated by LRG1 and AKT inhibitors at indicated levels, with DMSO as solvent controls. Long film exposures were done to show significant basal levels of P-AKT in CRC cells. **(C)** Western blotting showing P-eIF4B levels in CRC cells treated by LRG1 and p110β inhibitors GSK2636771 at indicated levels. Total HER3 (T-HER3), total eIF4B (T-eIF4B), total AKT (T-AKT), and β-actin are shown as loading controls.

**Figure S8. PI3K-PDK1-RSK inhibition, but not AKT inhibition, blocks eIF4B activation in HCT116 cells, related to** Figure 6. Western blotting showing P-AKT and P-eIF4B levels in HCT116 cells after treatments with LRG1 and indicated inhibitors. Total AKT (T-AKT), total eIF4B (T-eIF4B), and β-actin are shown as loading controls.

**Figure S9. LRG1 does not activate RSK2 but activate RSK3 in HCP-1 cells, related to** Figure 7. Western blotting showing P-RSK2 and P-RSK3 levels in HCP-1 cells treated by indicated peptides, CRC CM, or CM from EC-9 with control (siCTRL) or LRG1-specific siRNAs (siLRG1). P-RSK3 was not detected in other CRC cells, thus, not shown. Total RSK (pan RSK) and β-actin are shown as loading controls.

**Table S1. Lists of proteins identified by MS analysis of FPCL fractions of CRC and EC CM, related to Figure 1.** CRC and EC CM were processed by FPLC in parallel. Fractions 7 and 8 were combined for MS analysis. **(A)** proteins identified in CRC CM fractions. **(B)**, proteins identified in EC CM fractions. LRG1 in EC CM fractions were highlighted in red.

**Table S2. List of proteins identified by MS analysis of pull down assays with CM, related to Figure 1.** Grey, proteins pulled down by Ni-NTA control beads (Ctrl beads). Blue, proteins pulled down by HER3 ECD. Yellow, proteins pulled down by HER2 ECD. Columns Y—AO, direct comparison of each protein pulled down in indicated beads or ECD. LRG1 highlighted in red was uniquely found in HER3 ECD group. NF, not found.

**Table S3. Protein phosphorylation identified by global phospho-serine/threonine enrichment followed by LC-MS, related to Figure 4 and S5.** PDX-derived HCP-1 CRC cells were treated by CM from themselves (CRC CM) or EC-9 CM. Cell lysates were prepared from three independent experiments. **(A)** Results from 3 separate experiments with either CRC CM or EC CM treatments. **(B)** Results from 3 CRC CM- and 2 EC CM-treated samples after excluding one EC CM-tread sample that had low had low correlation coefficient with other two EC CM-treated samples. **(C)** Protein phosphorylation that are found in both groups and significantly increased in EC CM-treated samples. Ranked by log2 fold change. Key EIF components are highlighted in red. **(D)** Protein phosphorylation that are found in both groups and significantly increased in CRC CM-treated samples. Ranked by log2 fold change. **(E)** Changes derived from (C) and (D) and were used for IPA analysis. Ranked by phospho-log ratio. **(F)** Top EC CM-activated molecular and cellular functions determined by IPA analysis. Ranked by -log10 p value. Protein synthesis pathway is highlighted in red. **(G)** Top EC CM-activated canonical pathways determined by IPA analysis. Ranked by -log10 p value. EIF pathway is highlighted in red.

## MATERIALS AND METHODS

### Cell culture

Established human colorectal cancer (CRC) cell lines SW620 and HCT116 were purchased from American Type Culture Collection (ATCC, Manassas, VA). Murine CRC MC-38 cells were purchased from (Sigma-aldrich, St. Louis, MO). The patient-derived xenograft (PDX) derived CRC cell line HCP-1 was established and kindly shared by Dr. Lee Ellis from M.D. Anderson Cancer Center, as described previously ^14^. Primary human liver endothelial cells (EC-1, 3, 6, 8, 9) were isolated and established using MACS microbead-conjugated ant-CD31 antibodies and separation columns, as described previously ^12–14, 19, 51, 52^. All CRC cells were cultured in MEM (Sigma-aldrich) supplemented with 5% FBS (Atlanta Biologicals, Atlanta, GA), vitamins (1x), nonessential amino acids (1x), penicillin-streptomycin antibiotics (1x), sodium pyruvate (1x), and L-glutamine (1x), all from Thermo Fisher/Gibco (Grand Island, NY). Primary liver ECs were cultured in Endothelial Cell Growth Medium MV2 (PromoCell, Heidelberg, Germany) supplemented with 10% human serum (Sigma) and antibiotics-antimycotic (1x, Thermo Fisher/Gibco). All reagents and supplies use for cell culture can be found in key resources table.

HCP-1 cells and liver ECs were used within 10 passages, with approximately ∼1 week per massage. Mycoplasma contamination and short tandem repeat (STR) authentication tests for all cell lines are done every 6 months by the Genomics Core at Case Comprehensive Cancer Center, Case Western Reserve University. For primary cell lines (HCP-1 and liver ECs), genomic DNA samples from the original patient tissues were used for STR authentication.

### Animals

All experiments involving mice were approved by the Case Western Reserve University Institutional Animal Care Regulations and Use Committee (IACUC protocol # 2019-0087), and were done in 6-8 weeks old mice. Athymic nude mice and C57BL/6 mice were obtained from The Jackson Laboratory. Genetically engineered mouse model (GEMM) with systemic LRG1 knockout (LRG1 KO, C57BL/6-*Lrg1*^*tm1(KOMP)Vlcg*^/Mmucd) were created in prior studies^15^. *Lrg*^-/+^ heterogeneous breeding pairs were purchased from Mutant Mouse Resource & Research Centers (Stock number 048463-UCD), and were used for making LRG1 KO (*Lrg*^-/-^) and wild-type siblings (WT). Genotyping was done using primers, as described before ^15^: SU: TCCTGGTGGGAGAGGACTC. SD: GACCCCTGAAACAGACGTG. LacZRev: GTCTGTCCTAGCTTCCTCACTG. NeoFwd: TCATTCTCAGTATTGTTTTGCC.

### Reagents

The LRG1 monoclonal antibody 15C4 was developed in prior studies ^53^, and shared by Drs John Greenwood and Stephen E Moss at University College London, UK. Same levels of control human IgG antibodies were used from Invitrogen (Carlsbad, CA). HER3 extracellular domain (HER3 ECD, Cat# 10201-H08H), HER2 extracellular domain (HER2 ECD, Cat# 10004-H08H), recombinant human LRG1 (Cat# 13371-HCCH), FSTL1 (Cat# 10924-H08H), IGFBP4 (Cat# 10967-H08H), INHBA (Cat# 10429, -HNAH), and neuregulin-beta1 (Cat# 11609-HNCH, Ser 2-Lys 246 N-terminal fragment) were all from Sino Biological (Houston, TX). Human *NRG1*-specific siRNAs (5’-UCGGCUGCAGGUUCCAAAC, and 5’-GGCCAGCUUCUACAAGCAU), *ERBB3* (HER3)-specific siRNAs (5’-GCUGAGAACCAAUACCAGA and 5’-CCAAGGGCCCAAUCUACAA were used for gene knockdown), and a validated scrambled control siRNA were obtained from Sigma-Aldrich (St. Louis, MO). Human siRNA smart pools for *LRG1* (Cat# sc-97202), *RPS6KA1* (RSK1, Cat# sc-29475), *RPS6KA2* (RSK2, Cat# sc-36443), and *RPS6KA3* (RSK3, Cat# sc-36441) were obtained from Santa Cruz Biotechnology. Human CRC tissue microarrays were obtained from US Biomax (Cat# CO246, Derwood, MD). Inhibitors for AKT (MK-2206, Cat. #S1078), RSK (BI-D1870, Cat. #S2843), PDK1 (BX-795, Cat. #S1274), p110α (BYL719 Cat. #S2814), IRS1/2 (NT157 Cat. #S8228), eIF4 (4EGI-1, Cat. #S7369) were obtained from Selleckchem (Houston, TX). p110β inhibitor (GSK2636771 Cat. #HY-15245) was obtained from MedChemExpress (Monomouth Junction, NJ). Matrigel was from the R&D systems (Cat. # 3533-001-02. Minneapolis, MN).

### Conditioned medium (CM)

0.3×10^6^ CRC cells or ECs were seeded in T25 culture flasks overnight. Next day cells were washed two times with 1X PBS and then cultured in growth medium with 3ml 1% FBS (0.1×10^6^ cells/ml) for 48 hours. CM was harvested and centrifuged at 2,000 *g* for 5 minutes to remove cell debris. CM from each CRC cell line were used as controls.

### siRNA transfection

For each transfection, 1×10^6^ CRC cells were transiently transfected with 400 pmol siRNAs via electroporation by Neon Transfection System (Invitrogen, Carlsbad, CA) with 3 pulses of 10 msec at 1,600 V according to the manufacturer’s instructions. Cells were recovered in 5% FBS for 24-48 hours, cultured in 1% FBS overnight, and then subjected to CM for 30 minutes for Western Blotting, or 72 hours for the MTT assay. Transfection of ECs with *LRG1* siRNAs was carried out using Lipofectamine RNAiMAX (Life Technologies, CA), at a final concentration of 100 nM. Unless stated otherwise, cells were recovered for three days after transfection before used. Individual *NRG1*-specific siRNAs and 1:1 mixture of the two HER3 siRNAs were used. siRNA smart pools were used for other gene targets.

### Size execution fractionation

5mL 10x concentrated CM from ECs and CRC were injected into the Superdex 200 increase 10/300 GL size exclusion chromatography column (Sigma Cat. # GE28-9909-44). 200μl CM before column exclusion were saved as input for future analysis. After samples were loaded into the column, the first elution was performed with 5% of Hepes buffer (20mM Hepes, pH 7.5, 1M NaCl) for 5mL to remove weakly bound proteins. 20 fractions were collected with 0.5 mL/fraction. All fractions were stored at 4°C and used within 3 days, either for CRC cell treatments, Western blotting, or for mass spectrometry (MS).

### Pull-down Assay with HER3-ECD

2 μg purified HER3-ECD or control HER2-ECD (both have His-tag fused at the C-terminal) were added to 500 μl of 10X concentrated EC CM and incubated overnight at 4°C and with gentle rocking. Next day, 50 μl Ni-NTA beads were added to the mixture and incubated for 1 h at 4°C with gentle rocking, and then centrifuged at 13,000 *g* for 2 minutes. After removing the supernatant, Ni-NTA beads were wash twice with 0.5 ml 20mM imidazole, and then incubated in 50 μl of 500 mM imidazole for 5 minutes at room temperature, and centrifuged at 13,000 *g* for 2 minutes. Supernatant with eluted soluble proteins were used either for treating CRC cells, Western blotting, or for MS analysis. Ni-NTA beads were also used for Western blotting.

### Mass spectrometry (MS)

MS analysis for fractionated CM or for pulled-down proteins by HER3-or HER2-CD were all done at Taplin Mass spectrometry Facility, Harvard Medical School, Boston, MA. Samples were treated with trypsin overnight at 37°C and acidified by spiking in 20ul 20% formic acid solution and then desalted by STAGE tip. Samples were then reconstituted in 10 µl HPLC solvent A. After equilibrating the column, each sample was loaded via a Famos auto sampler (LC Packings, San Francisco CA) onto the column. A gradient was formed and peptides were eluted with increasing concentrations of solvent B (97.5% acetonitrile, 0.1% formic acid). Eluted peptides were subjected to electrospray ionization and then entered into an LTQ Orbitrap Velos Elite ion-trap mass spectrometer (Thermo Fisher Scientific, Waltham, MA). Peptides were detected, isolated, and fragmented to produce a tandem mass spectrum of specific fragment ions for each peptide. Peptide sequences (and hence protein identity) were determined by matching protein databases with the acquired fragmentation pattern by Sequest (Thermo Fisher Scientific).

### Western blotting

CRC cells were treated with indicated treatments for 30 minutes. Cell lysates were processed and run through SDS-PAGE gel electrophoresis as described previously^13, 14, 19, 51, 52^. For CM samples and or bead-bound proteins in pull-down assays, CM and beads were boiled with SDS-PAGE buffer for 5 minutes. HRP conjugated β-actin antibody was from Santa Cruz Biotechnology, and LRG1 antibody was from R&D Systems (Cat# AF78900). All other antibodies were from Cell Signaling (Beverly, MA). For each experiment, protein samples were loaded to two gels and processed at the same time for separately probing for antibodies specific to phosphorylated proteins and total proteins, and for detecting proteins that have similar sizes. All membranes were probed with β-actin as loading control and a representative image was shown for each experiment. Each Western Blotting figure shows representative results of at least three independent experiments.

### Biolayer Interferometry (BLI) assay

HER3-ECD, human recombinant LRG1 and neuregulins were purchased from Sino Biological and reconstituted according to manufacturer instructions. The binding affinity between HER3-ECD and LRG1 or neuregulins was measured by the Octet RED96 system (Sartorius), at 25 °C in 1X PBS buffer. Ni-NTA sensors were pre-equilibrated in PBS for at least 10 min before use in experiments. HER3-ECD with His-tag was loaded onto the Ni-NTA biosensors for 180 seconds. For multiple cycle kinetics, six sensors were immersed with LRG1 or neuregulins with concentrations in the range from 4 nM to 1 µM for 120 seconds for the association step, with PBS buffer as a control. Sensors were then immersed in PBS buffer for 240 seconds for the dissociation step. The binding kinetics Kd was determined by the 1:1 Langmuir interaction model by analyzing the data with the Octet Analysis Studio 12.2.2.26.

### Phospho-proteomics analysis

HCP-1 CRC cells were seeded at 5 × 10^6^ cells/well in 10 cm plates, cultured in 1% FBS overnight and then treated with EC-9 CM for 30min, with HCP-1 CM as controls. Cells were scrapped off plates on ice in 1x cold PBS and washed twice with PBS. Cell pellets from three independent experiment were sent to the Proteomic and Metabolomics core at Lerner Research institute, Cleveland clinic, Cleveland, OH, for global serine (S) and threonine (T) protein phosphorylation analysis. Cell pellets were digested by trypsin for cleavage after Lys/Arg residues, including Lys/Arg-Pro bonds, and then subjected to phospho-S/T enrichment using Fe-NTA spin columns from Thermo Scientific. The enriched samples were then analyzed by LC-MS/MS by a Thermo Scientific Fusion Lumos MS system.

Phospho-peptides were identified by comparing all of the experimental peptide MS/MS spectra against the UniProt human proteome database downloaded on April 19, 2018 using an Andromeda search engine (PMID: 21254760) integrated into the MaxQuant version 1.6.3.3 (PMID: 19029910). Carbamidomethylation of cysteine was set as a fixed modification, whereas variable changes included phosphorylation of serine/threonine/tyrosine, oxidation of methionine to methionine sulfoxide and acetylation of N-terminal amino groups. For trypsin specificity, the minimum peptide length was set to 7, the maximum missed cleavage was set to 2, and the cutoff false discovery rate (FDR) was set to 0.01. Match between runs (match time window: 0.7 min; alignment time window: 20 min) was enabled. For computation for principal component analysis and volcano plotting, identified peptides were filtered with S0=0.1 to minimize the coefficient of variation for computation^54^. Remaining options were kept as default. Perseus tool (PMID: 27348712) was used to quantify the identified phosphopeptides. The “Phospho(STY)Site.txt” file generated by the database search was incorporated into Perseus, then the phosphorylation sites identified were filtered for those that were confidently localized (localization probability > 0.75), and Log2 transformed to normalize the data distributions. Student’s two-sample t-test was used to estimate the significance levels for the difference in the phosphorylation levels at individual sites between CRC CM-treated and EC CM-treated cells. Greater than 2-fold difference in phosphorylation level with a p-value of <0.05 was considered significant. QIAGEN Ingenuity Pathway Analysis (IPA) was used for pathway and functional analyses of identified significantly and differentially phosphorylated proteins in HCP-1 cells treated by control HCP-1 CM or EC CM with imputed ± 6.64 fold changes for unique changes.

### MTT assay

CRC cells were seeded at 3,000 cells/well in 96-well plates, cultured in 1% FBS overnight and then treated with CM for 72 hours. When the LRG1 antibody 15C4 was used, cells were pretreated with 500 µg/ml 15C4 in 1% FBS medium for 6 hours, and then cultured in CM with or without 15C4 for 72 hours. Cell viability was assessed by adding MTT substrate (0.25% in PBS, Sigma) in growth medium (1:5 dilution) for 1 hour at 37 °C. Cells were washed with 1x PBS and then incubated in 50 µl DMSO. Optical density was measured at 570 nm and relative MTT was presented as % of control.

### LRG1 depletion from CM

ECs were transfected by control or *LRG1-*specific siRNAs and recovered for 72h. CM from transfected cells were then prepared as described before. For LRG1 immuno-depletion by antibodies, 5 µg control IgG or anti-LRG1 antibodies (both from R&D Systems) were added to 500 μl of 3X CM (1000 μg total proteins) from un-modified ECs After rocking overnight at 4^°^C, a 20 μl mixture of protein A/G PLUS-Agarose beads was added to CM and rocked at 4°C for 1 hour. CMs after immuno-depletion were used for treating CRC cells and tumors, or Western blotting. Beads with proteins were also used for Western blotting.

### Global Protein Synthesis assay

The Global protein synthesis assay kit from Abcam (Cat# ab273286, Waltham, MA) was used following to the manufacturer’s instructions. In brief, CRC cells were seeded 100,000 cells/well in 96-well plates, cultured in 1% FBS overnight and then incubated with CM and O-Propargyl-puromycin (OP-puro) analogs for *de novo* protein labeling for 24 hr. Next, day, cells were fixed, permeabilized, labeled with florescence probes. Cells were also labeled with DAPI for total loading controls. Intensities of florescence probes and DAPI were measured by an EVOS cell imaging system (Invitrogen), and presented as % of control relative to DAPI.

### Subcutaneous (subQ) xenograft model

1×10^6^ HCP-1 or 2×10^6^ SW620 cells were injected subQ into the right flanks of athymic nude mice in 100 μL growth-factor-reduced Matrigel. After xenografts were confirmed, mice were randomized and then treated with 3x concentrated CRC or EC CM once a week by injecting CM subQ into the spaces between xenografts and skin tissues. Tumor volumes were measured by a caliper. All mice were euthanized when 3 mice from any group became moribund or tumor sizes reach 1000 mm^3^, and tumor tissues were harvested. For 15C4 treatments, mice were treated with CM and with either IgG control antibody or 15C4 20 mg/kg twice a week by intraperitoneal (IP) injection.

### Orthotropic allograft mCRC model

Luciferase reporter-labeled MC-38 murine CRC cells were suspended in an inoculation matrix (1:1 mix of growth-factor-reduced Matrigel and 1x PBS) and injected 1×10^5^ cells in 30 µl/injection into the liver of C57BL/6 mice. For studies in GEMM with LRG1 knockout, MC-38 cells were injected into mice with systemic knockout of LRG1 and wild-type siblings. For studies with 15C4 treatments, wild-type C57BL/6 mice were used. After implanting MC-38 cells, tumor burden was assessed weekly by bioluminescence with the *In Vivo* Imaging System (IVIS) and D-Luciferin substrate (Xenogen, Alameda, CA) according to the manufacturer’s instructions. When tumor burden was confirmed by IVIS on Day 28 after implant, mice were randomized and then treated with control IgG or 15C4 antibodies (20 mg/kg by IP twice a week). Mice were sacrificed when became moribund. CRC tumors in the liver were collected for analysis. Mice overall survival were determined by time to death with Kaplan-Meier plots. Tumor burden over time of each mouse was presented in spider plots. Estimated mean tumor burden over time was analyzed using a longitudinal mixed effect linear regression model. The model included fixed effects for time and group, and random effects to account for patient correlation. The correlation structure was modeled using an autoregressive (AR1) structure to account for the repeated measures. Tumor size was log-transformed prior to analysis to linearize the growth pattern and stabilize the variance. Mean log-tumor sizes and their corresponding 95% confidence intervals were estimated at each time point for both groups. These were subsequently back-transformed and plotted to visualize tumor growth patterns and compare between the treatment vs control groups. *t*-test was adopted to test the difference of tumor size between case and control group at timepoints 28, 35, 42, 49, 56 days. The analyses were conducted using the Linear and Nonlinear Mixed Effects Model (NLME) package in R.

### Immunohistochemistry (IHC)

Tumor tissues were fixed in 4% paraformaldehyde, dehydrated in ethanol, and embedded in paraffin. Sections were cut 10 μm in thickness. Human CRC tissue microarrays in 10 μm were purchased from TissueArray (Cat# CD702d). IHC staining was done with antibodies against P-eIF4B S422 (1:200; Cell Signaling Technology, Cat# 3591), P-HER3 Y1289 (1:100; Cell Signaling Technology, Cat# 2842), and Ki-67 (1:200, Cell Signaling Technology, Cat# 9449T). Isotype-matched immunoglobulin was used as a control. The HRP-conjugated anti-rabbit secondary antibody (Cell Signaling Technology, Cat# 8114p) were used with the DAB substrate kit (Vector Laboratories, Newark, CA, SK-4800) for staining. Stained sections were examined under a light microscope (Olympus BX51). Intensity of the staining was quantified by QuPath software (NIH) and presented as density relative to control groups with CRC CM.

### Statistical analysis

For *in vitro* assays, all quantitative data were reproduced in at least three independent experiments with multiple measures in each replicate. Groups were compared by unpaired two-tailed Student’s *t*-test and data was expressed as means -/+ standard error of the mean (SEM.) with significance of P<0.05. For *in vivo* assays, Wilcoxon rank-sum test was used for tumor volume and burden change over time, and for Kaplan-Meier plots for overall survival. One-way ANOVA was used for comparing between-group difference analysis after collecting tissues at the end of experiments, and tissue IHC analysis. *In vivo* data was expressed as means -/+ standard deviation (SD) with significance of P<0.05. For all figures, *p<0.05, **p<0.005, ***p<0.0005, ****p<0.00005 by indicated statistical test analysis.

## Notes

**Conflict of Interest** The authors declare no potential conflicts of interest.

### Competing Interest Statement

The authors have declared no competing interest.

### Summary of Updates

revised manuscript and updated figures

