## Supplemental Figures for "LRG1 is a novel HER3 ligand to promote growth in colorectal cancer"

### Figure S1

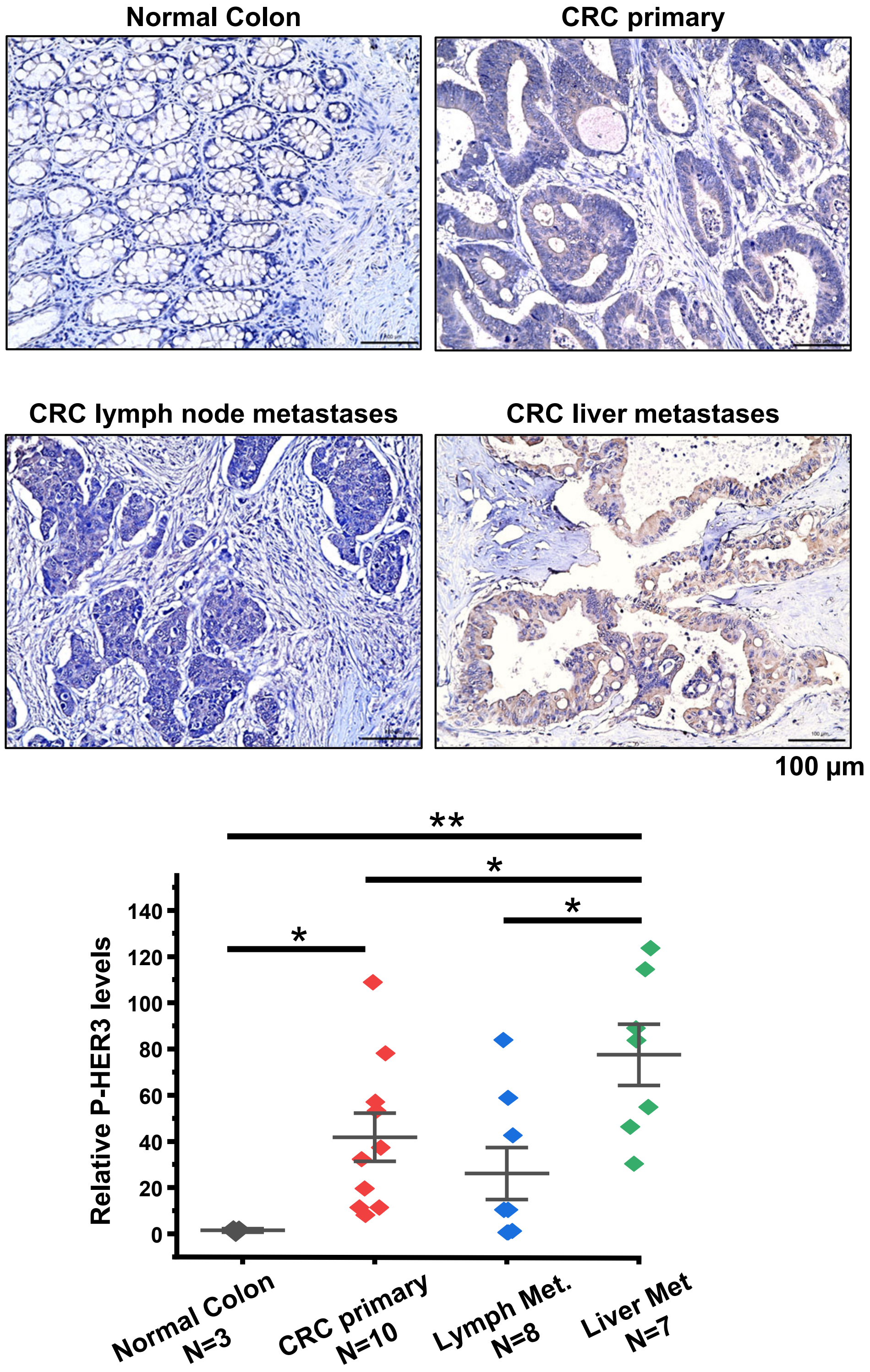

Normal Colon
CRC primary
CRC lymph node metastases
CRC liver metastases
100 µm
**
*
*
*
Relative P-HER3 levels
0
20
40
60
80
100
120
140
Normal Colon N=3
CRC primary N=10
Lymph Met. N=8
Liver Met N=7

### Figure S2

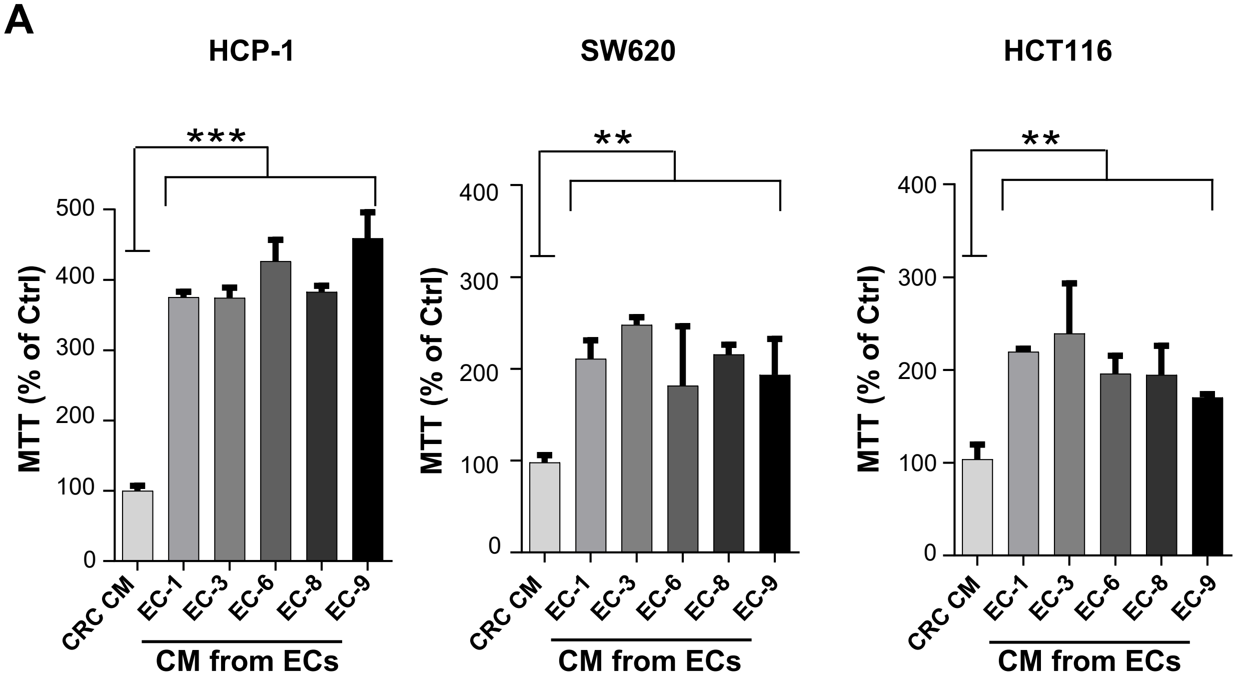
A
HCP-1
SW620
HCT116
***
**
**
MTT (% of Ctrl)
0
100
200
300
400
500
CRC CM
EC-1
EC-3
EC-6
EC-8
EC-9
CM from ECs

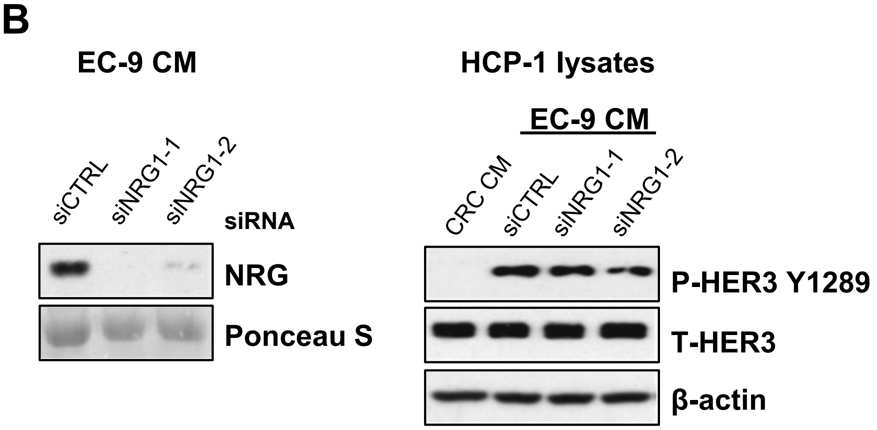
B
EC-9 CM
siCTRL
siNRG1-1
siNRG1-2
siRNA
NRG
Ponceau S
HCP-1 lysates
EC-9 CM
CRC CM
siCTRL
siNRG1-1
siNRG1-2
P-HER3 Y1289
T-HER3
β-actin

#### Figure S3, related to Figure 1

A

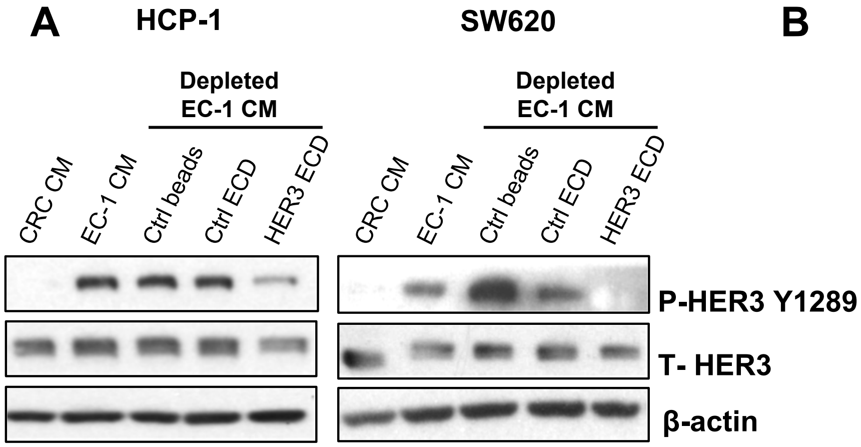


B

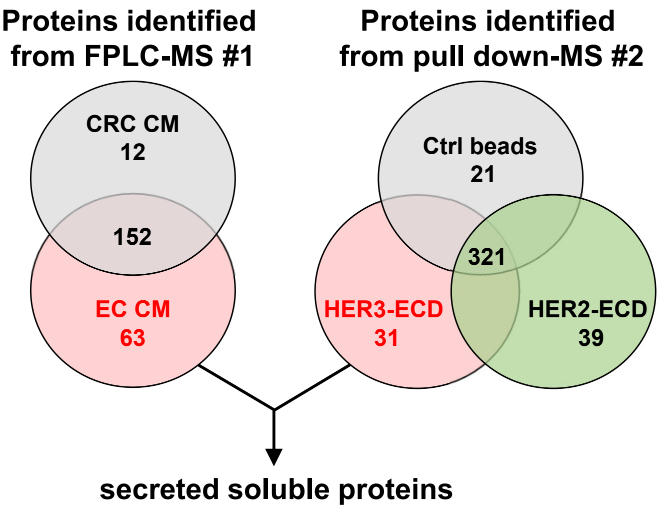


C

Top candidates

| ID | Intensity |
|---|---|
| FSTL1 | 1.40E+06 |
| IGFBP4 | 5.90E+05 |
| LRG1 | 5.60E+05 |
| INHBA | 4.00E+05 |
| NRP1 | 3.50E+05 |
| LTBP3 | 9.60E+04 |
| TGFB1 | 3.50E+04 |

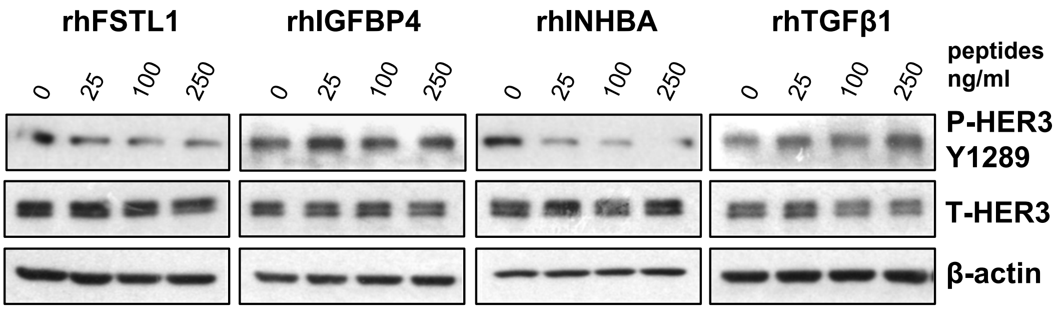


D

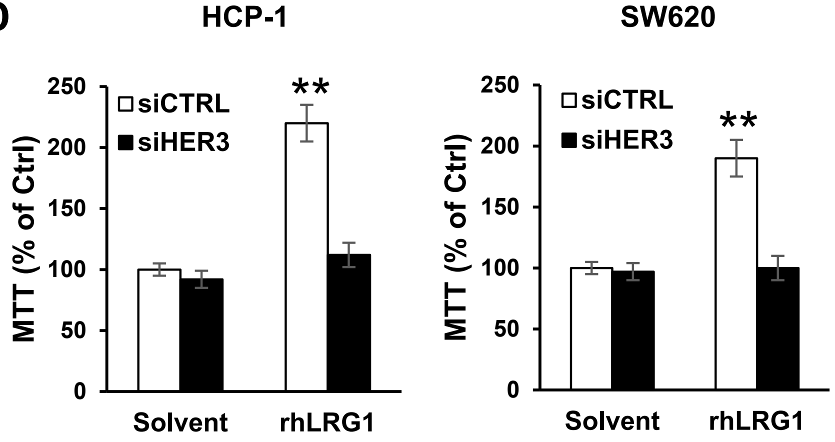


E

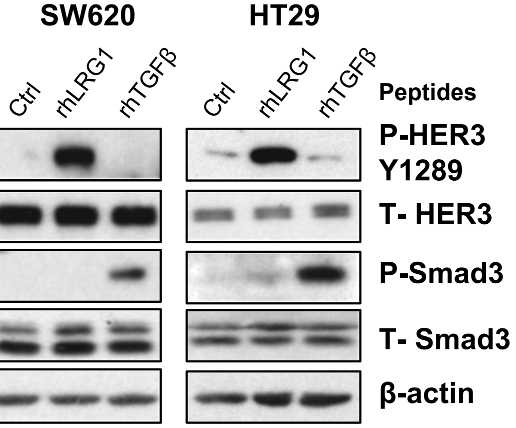


F

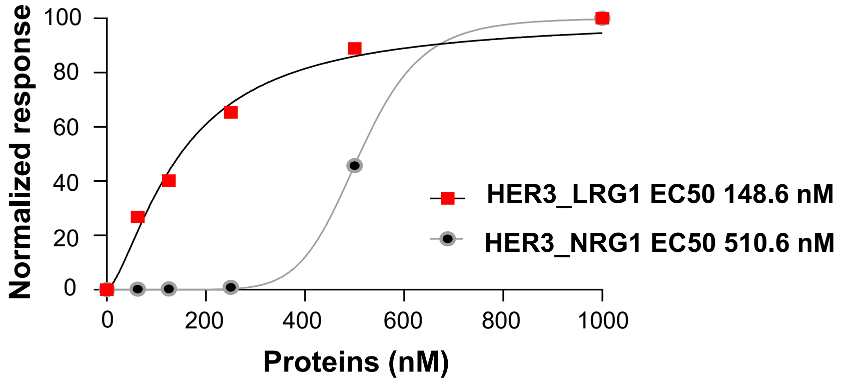

### Figure S4, related to Figure 3

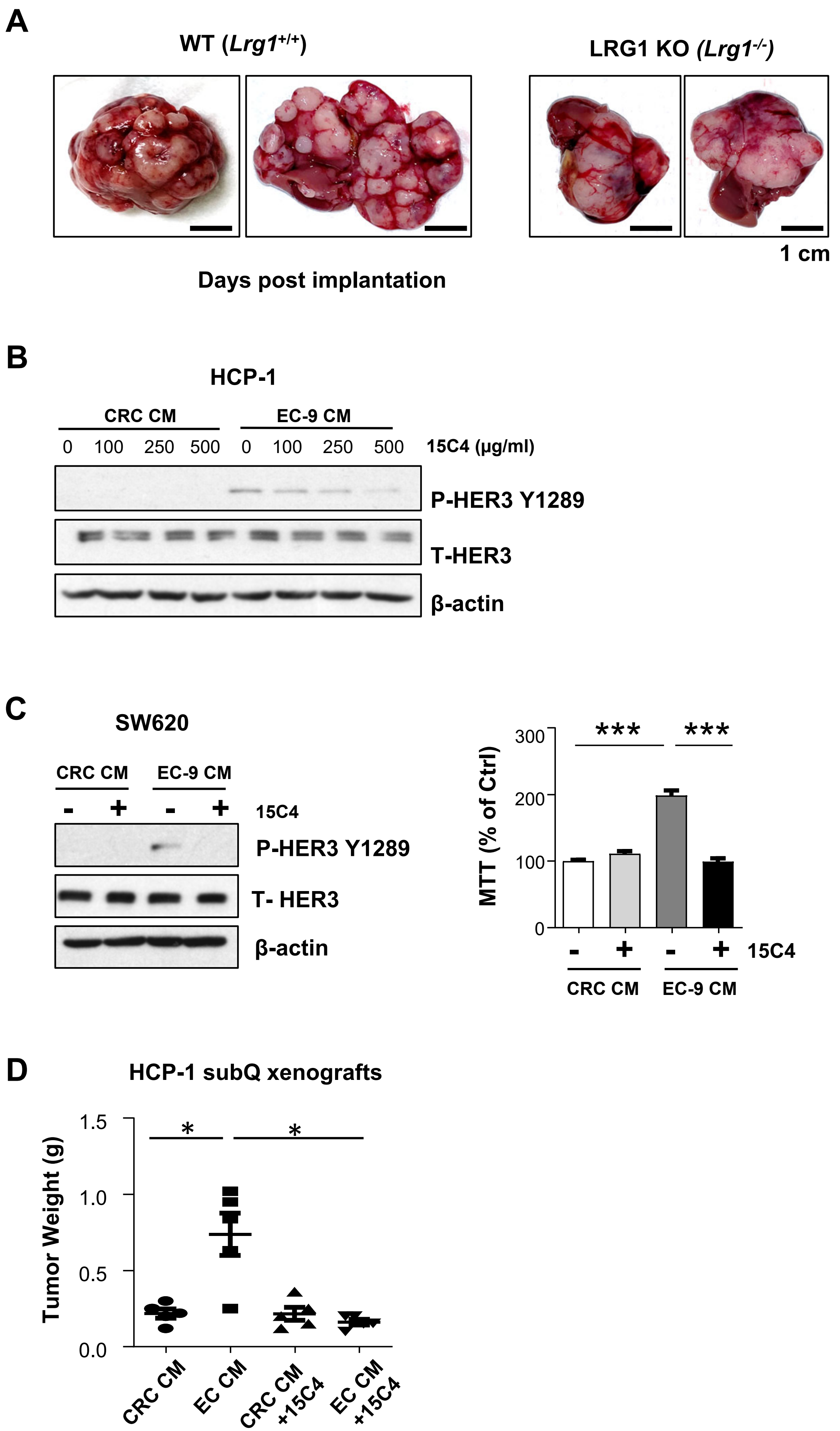

**Figure S5, related to Figure 4**

A

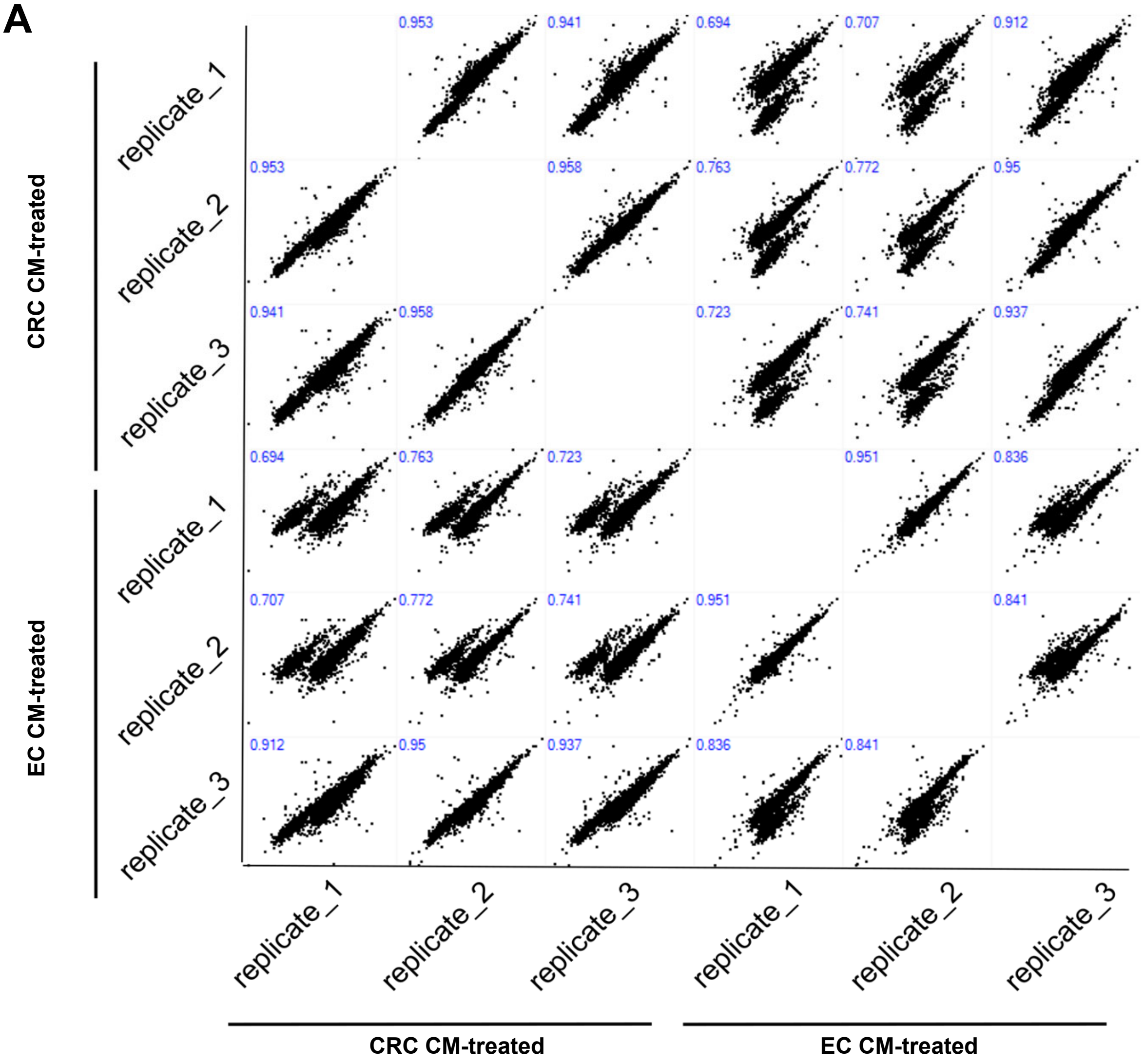


B

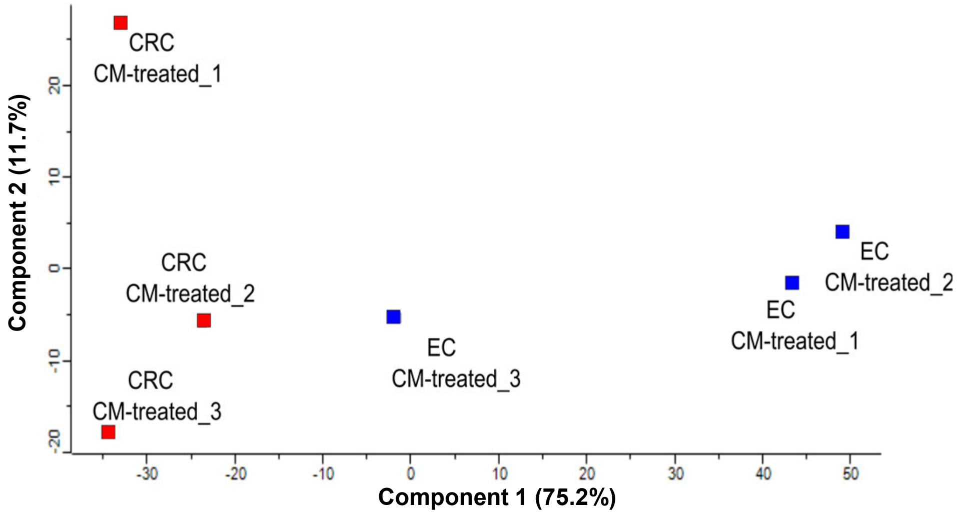

#### Figure S6, related to Figure 5

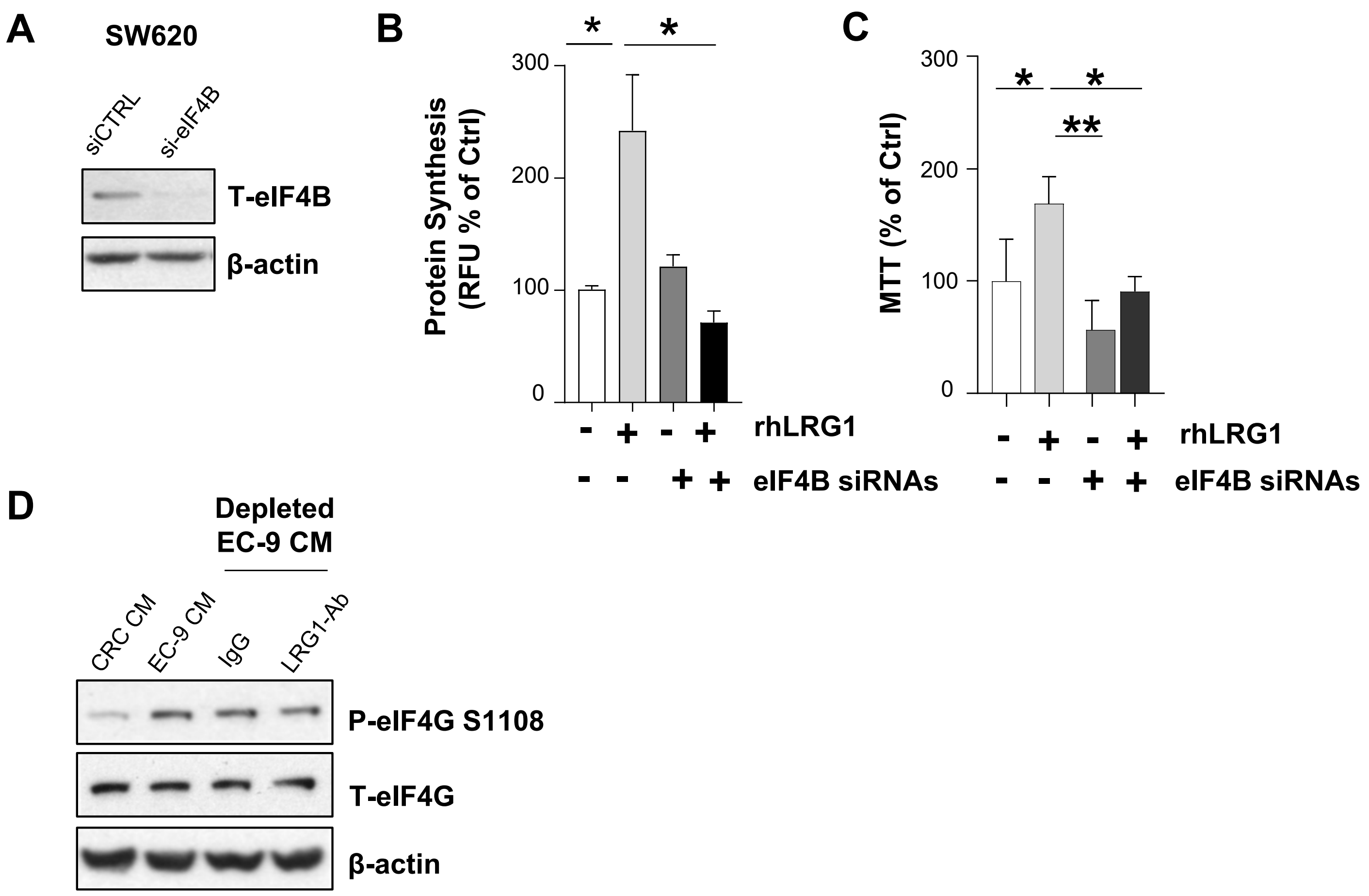

#### Figure S7, related to Figure 6

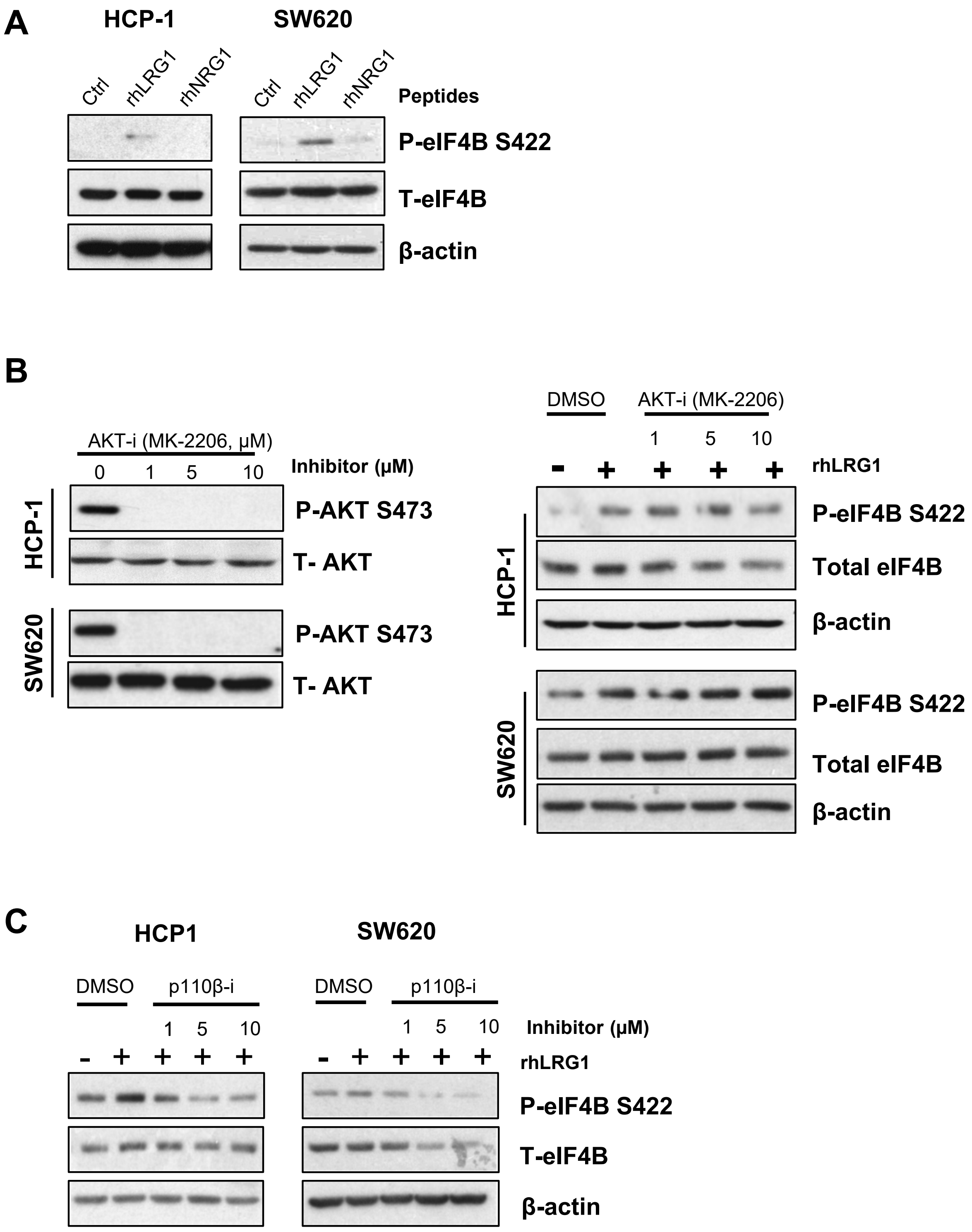

### Figure S8, related to Figure 6

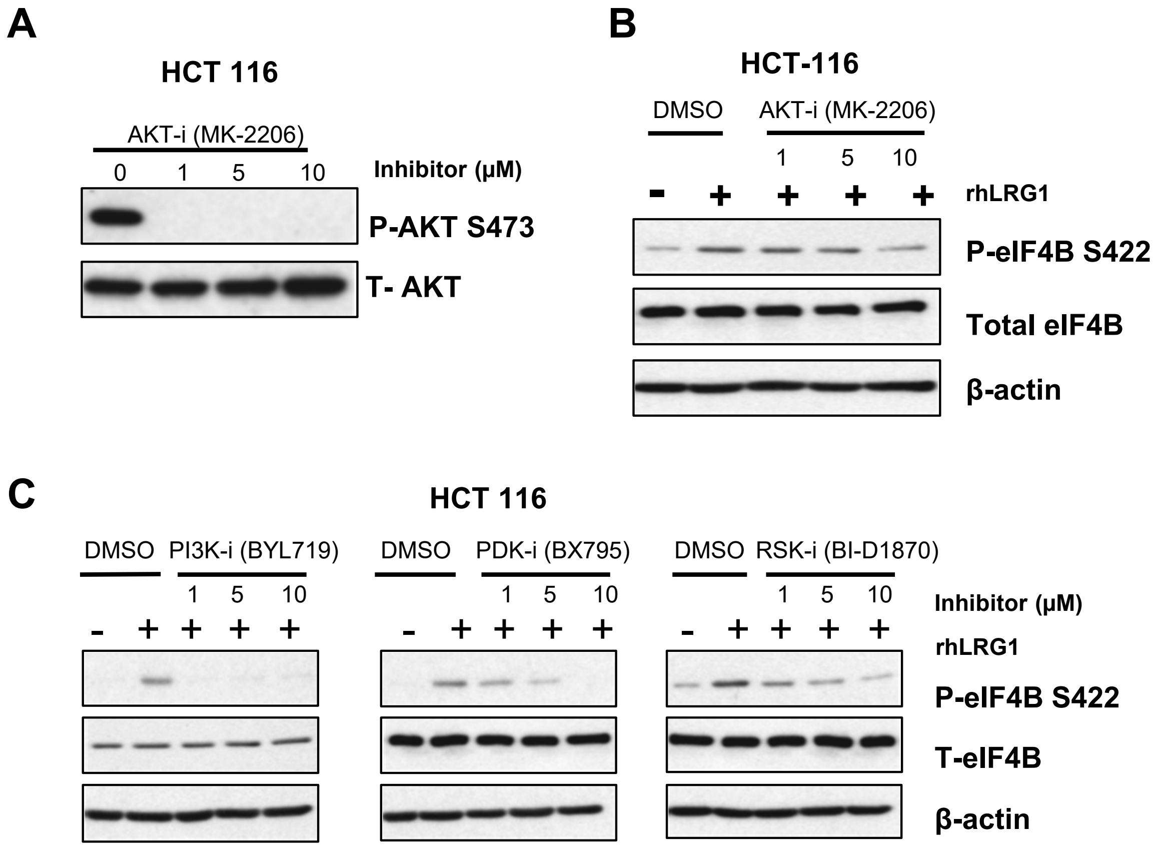

### Figure S9, related to Figure 7

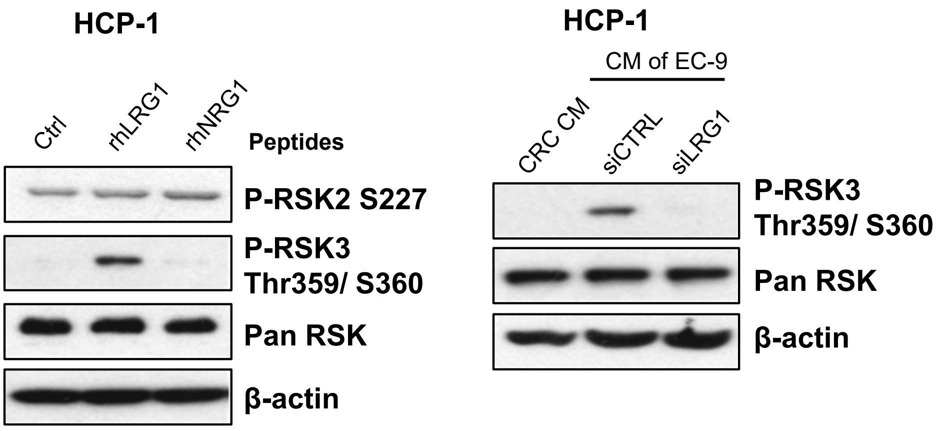
